# Investigating active dynamics of contractile actomyosin gels with Micro Particle Image Velocimetry (Micro-PIV) analysis

**DOI:** 10.1101/2025.08.14.670336

**Authors:** Sakshi Choudhary, Subhaya Bose, Yuval Amit, Daniel Sevilla Sanchez, Gefen Livne, Kinjal Dasbiswas, Anne Bernheim-Groswasser

## Abstract

Micro Particle Image Velocimetry (Micro-PIV), an advanced imaging technique, enables high-resolution velocity field measurements by tracking fluorescent tracers in microscopic environments. Here, we adapt conventional micro-PIV to study the rapidly contractile dynamics of active poroelastic gels. We demonstrate how frame-to-frame correlation improves signal-to-noise ratios and how the elastic nature of the solid phase of the gel can be included in the analysis. To do this, we average the gel displacement data under an axisymmetric assumption to extract radial strain profiles that reliably reveal local deformations of the gel. By analyzing gels of varying shapes, we further show that our method extends robustly to gels that are not completely circular or that do not displace symmetrically towards their geometric center. The analysis reveals common underlying features in the radial profiles of gel deformation. These strain profiles will allow the inference of the spatial and orientational distribution of motor-generated active stresses with appropriate constitutive models for the gel mechanics. Our findings emphasize the importance of tailored micro-PIV methodologies for analyzing complex fluids, particularly autonomously contracting poroelastic materials. This approach significantly enhances understanding of cytoskeletal dynamics and self-organization processes, with broad implications for cell motility, morphogenesis, and active matter physics.

## Introduction

Cell shape changes are essential for biological processes such as cell division and cell motility during embryonic development, wound healing and immune responses (1). In animal cells, such shape changes are typically driven by the dynamic reorganization of their actomyosin cytoskeleton, a network of actin filaments, myosin molecular motors, and accessory proteins. Myosin motors consume ATP to slide actin filaments according to their polarity (2), leading to active contractility and self-organization of the cytoskeletal network (3). These active and chemo-responsive mechanical forces enable cellular functions such as shape changes and movement (4,5), and are an example of active, adaptive gels that exhibit both viscous and elastic response (6–8).

However, the complexity of the cytoskeleton, with its interconnected chemical and mechanical processes spanning diverse spatial and temporal scales, poses significant challenges for the complete understanding of its structural changes during cell motility and morphogenesis. In vitro reconstituted systems allow for controllable investigations using a minimal set of components, actin filaments, myosin II motors, ATP, and an actin crosslinker (such as fascin), which together form contractile elastic networks (9–12). These actomyosin gels, which mimic the mechanical properties of the cellular cytoskeleton, provide valuable insight into active force generation and contractility observed in processes such as shape changes, motility, and tissue mechanics. Myosin motor-induced active forces have been shown to drive complex 3D shape changes in active gel sheets (13,14), and identifying the spatiotemporal distribution of these forces requires tracking the gel dynamics. Notably, these dynamics exhibit poroelastic effects, where motor-driven contraction of the elastic actin network leads to the outflow of interstitial fluid (13), consistent with observations from single cell rheology of the cell cytoplasm (15). Consequently, probing such complex two-phase dynamics requires modifications to conventional methods of tracking microscale flows.

Particle Image Velocimetry (PIV), Traction Force Microscopy (TFM), Optical Flow, and Micro Particle Image Velocimetry (Micro-PIV) represent cutting-edge experimental techniques at the forefront of fluid mechanics research (16–18). Particle Image Velocimetry (PIV) is a technique commonly used to analyse the velocity field within a fluid flow on macroscopic scales (19,20). In a typical PIV experiment, the fluid is seeded with tracer particles that are sufficiently small to closely follow the flow but large enough to scatter enough light for detection on standard imaging systems. A laser sheet illuminates a thin slice of the flow, and high-speed cameras record successive images. By using cross-correlation algorithms on these image pairs, the displacement of the material elements, and thus the displacement vectors, are calculated over a relatively large area. This method is highly valued for its non-intrusive nature and its ability to provide whole-field measurements rather than single-point data. Traditional PIV employs relatively large tracer particles ranging in size from tens to hundreds of micrometers (19,20), though smaller particles ranging from sub-micrometers to micrometers length scales have been employed for analyzing the solvent flow in contractile poroelastic cytoskeletal networks (13). PIV offers high spatial resolution for macroscopic flow fields and is frequently used in applications such as aerodynamics, environmental fluid dynamics, and industrial turbulent flow research. PIV requires only standard cameras and image equipment. Data analysis entails processing images to capture macroscopic flow patterns, making it perfect for investigating large-scale flow behaviour.

Traction Force Microscopy (TFM) is commonly used to estimate the mechanical forces that cells exert on soft substrates (17). This method involves embedding fluorescent beads within a deformable material and tracking their displacement as the substrate responds to cellular forces. The resulting deformation field is then used to reconstruct traction stresses through computational modeling. High-resolution implementations have improved the sensitivity and accuracy of TFM, as shown in the work of Sabass et al. (2008) (17). However, the approach requires detailed knowledge of the substrate’s mechanical properties and involves solving an inverse problem, which can introduce uncertainty in complex or dynamic systems. Optical flow presents another method for motion estimation, using changes in pixel intensity across image sequences to calculate displacement fields. It is widely used in biological imaging due to its speed and simplicity, particularly for analyzing movements such as cell edge dynamics or tissue deformation (21,22). Despite its advantages, optical flow is sensitive to image quality and may not perform well in systems with non-uniform contrast or irregular motion.

In comparison, Micro-Particle Image Velocimetry (Micro-PIV) directly measures displacement by tracking fluorescent tracer particles under high magnification. This technique does not rely on mechanical modeling or pixel intensity changes, making it a more reliable choice for quantifying deformation in soft, contractile biological materials like actomyosin gels. Unlike traditional PIV, which operates on millimeter to meter scales (19,20,23), Micro-PIV is tailored to investigate fluid flow phenomena at microscopic scales, typically within microfluidic channels or devices (24–26). Due to the reduced dimensions, specialized optics, such as high-magnification microscope objectives, are required to resolve the tracer particles (usually of micron-sized scale) and finer flow details. Often the technique employs fluorescence illumination to improve particle contrast against the background, and due to the high magnification, issues like depth-of-field are more pronounced, which can lead to challenges in ensuring that only particles within a very narrow slice are imaged accurately.

Micro-PIV methodology addresses the limitations of conventional PIV when applied to micron-sized systems, where reduced dimensions and low flow rates demand high precision. At these scales, noise and low particle density can significantly hinder accurate velocity measurement, making ensemble averaging of multiple image pairs essential for extracting statistically reliable flow fields. One of the distinguishing features of Micro-PIV is its superior spatial resolution, which enables detailed investigation of fluid behavior in microchannels and small-scale systems, both artificial and native (e.g., biological). In particular, it allows for the visualization of flow patterns, identification of vortices, and measurement of velocity gradients within microchannels and complex microfluidic networks (18), which is especially important in applications such as lab-on-a-chip devices. Micro-PIV is particularly well-suited for studying flows at low Reynolds numbers, common in microfluidic systems and Micro-Electro-Mechanical Systems (MEMS) (27). Understanding these flows is crucial not only for the development and optimization of microscale technologies but also for resolving the dynamics of complex biological systems, which remain poorly understood.

Micro-PIV finds wide-ranging applications across various disciplines. In microfluidics, it helps elucidate fundamental fluid behavior within confined geometries, facilitating the optimization of microfluidic devices for applications such as lab-on-a-chip systems, chemical synthesis, and biological analysis (24,25,28). In biomedical engineering, Micro-PIV contributes to understanding blood flow dynamics in micro vessels, aiding in the development of targeted drug delivery systems and diagnostic tools (18,29–31). Additionally, in MEMS (Micro-Electro-Mechanical Systems), Micro-PIV assists in optimizing fluidic actuation and cooling systems within miniature devices (27). Overall, Micro-PIV is a powerful tool for studying fluid dynamics at the microscale, providing invaluable insights into the behaviour of fluids in complex microenvironments. Its high-resolution measurements and non-invasive nature make it indispensable for advancing research and innovation in microfluidics, biomedical engineering, and MEMS technologies.

In this study, Micro-PIV was employed to analyse the contraction dynamics of actomyosin gels of controlled composition and elastic properties. We adapt conventional micro-PIV to a system with poroelastic, contractile dynamics that dramatically reduces in size during the period of observation. Instead of placing invasive beads, we track the local inhomogeneities in gel density, caused by aggregation of actin by myosin (actin foci) that appear as bright spots in network fluorescence signal which we utilize for PIV analysis (32,33). Unlike a purely viscous fluid, the spontaneous generation of complex and stable final 3D shapes from an initially flat geometry suggest that these gels undergo buckling due to imbalance in in-plane elastic stresses (14,34). This manuscript describes the principles, challenges, and methodology for the adaptation of micro-PIV to analyze the complex dynamics in the actomyosin gels across varying scales.

## Methods

### Micro Particle Image Velocimetry (Micro-PIV) of contractile poroelastic actomyosin gels

Here we extend this methodology to explore the material and flow properties of contractile actomyosin gels of various shapes. The actomyosin networks are formed upon mixing 5μM actin monomers, 16.7 nM myosin motors - added in the form of large filaments - each consisting of ∼150 myosin dimers, 280 nM of the passive crosslinker fascin, and 2mM ATP and an ATP regenerating system which maintains the level of ATP constant in the solution. A drop of this solution is squeezed between two PEG-passivated glass coverslips, where the drop volume and the spacing between the two coverslips set the initial actomyosin gel size (13,35). Network formation initiates by actin filaments polymerization, and their crosslinking by fascin and myosin motors, to form an interconnected active network.

Myosin motors generate contractile forces by interacting with the actin filaments in an ATP-dependent manner. This interaction produces active stresses, leading to gel contraction, which can be followed by labelling the actin monomers (or myosin motors) using fluorescent markers (13,35). Figure 1 outlines the step-by-step process of Micro-PIV analysis used for the analysis of such a contracting actomyosin gel of disc of initial radius *R*. The process starts with the acquisition of raw images via 2d fluorescence imaging using high-speed cameras (13), capturing the movement of the previously described localized fluorescent actin foci in the gel (Fig. 1a). These images are then optimized through background subtraction to remove noise and, if necessary, stabilized to correct for any translational drift, in sample x and/or y position (Fig. 1b). Next, the optimized images undergo masking, where specific regions of interest (ROIs) are selected for analysis. Here, the ROI is the actomyosin gel (Fig. 1c), excluding irrelevant areas such as the outer background solution and drop edge, the white circle in Figs. 1a and b. The displacement vectors of the “material elements” are then calculated using ensemble correlation through Micro-PIV (Fig. 1d), providing velocity measurements by analysing the displacement between successive image pairs (26). Finally, strain analysis is performed, leveraging the displacement vectors to calculate strains within the gel, offering insights into the deformation and local dynamics of the gel (Fig. 1e).

**Figure 1.**
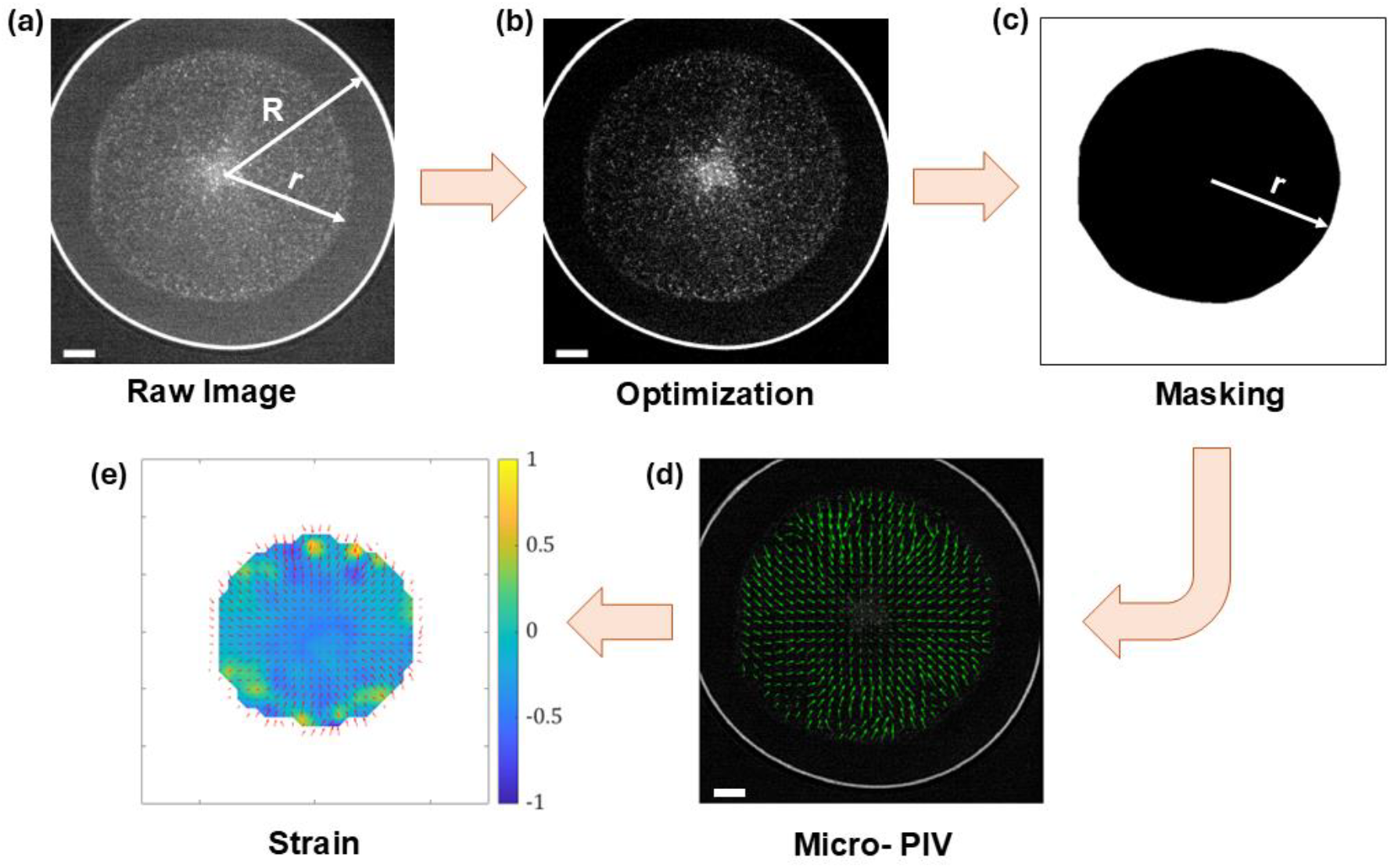
Flowchart describing the overall process of the micro-PIV analysis of contractile actomyosin gels. (a) Starting from the acquisition of the raw images (Here, actin is labelled fluorescently with Alexa-Fluor 488), (b) Optimization, which involves background subtraction and image stabilization (if needed), (c) Masking of the optimized images for the region of interest (here, the actomyosin gel), (d) Displacement vectors through ensemble correlation by using Micro-PIV and, (e) Strain analysis. *R* (= 1600μm) and *r* are the initial and temporal gel disc radius. Bars are 300μm.

#### Displacement of Particles

In PIV, the displacement of material elements between two consecutive image frames is an important characteristic for estimating the material velocity. The displacement is denoted as *Δ****x***. This displacement of the *i*^*th*^ material element can be expressed as the difference in their positions at two distinct time instances:

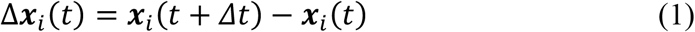

where ***x***_***i***_(*t*) and ***x***_***i***_(*t* + *Δt*) are the positions of the *i*^*th*^ material element at time *t* and *t* + *Δt*, respectively, such that Δ***x***_***i***_(*t*) represents the displacement vector between the two time-points separated by Δ*t*. Note that we use bold face symbols to indicate vector quantities.

#### Velocity Estimation

The velocity of the *i*^*th*^ material element can thus be estimated from the displacement over the time interval Δ*t*, and is given by

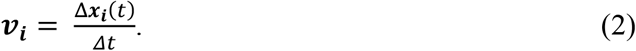

By interpolating this Lagrangian information for the displacement of individual material elements, we can construct the Eulerian displacement or velocity field, ***v***(*x, t*). Equation 2 represents the velocity (***v***), computed as the displacement divided by the time interval. Choosing an appropriate *Δt* is essential for balancing the trade-off between measurable displacement and maintaining a strong correlation. For accurate velocity measurements, *Δt* should be chosen so that the displacement of particles is large enough to be measured accurately but not so large that the particles leave the interrogation window, a small sub-region in the image used for calculating tracer particles’ displacement.

In PIV measurements, the choice of time interval *Δt* significantly affects both temporal resolution and velocity accuracy. A smaller *Δt* may be necessary to capture high-speed flows, but it can also lead to issues like particle overlapping or blurring, which complicate tracking. To accurately extract velocity fields, multiple particles must be reliably tracked across consecutive image frames, requiring a high seeding density.

- **Small** *Δt*: If *Δt* is too small, particles may not move enough between frames, resulting in displacements that are too small to measure precisely. In such cases, the correlation peak can be broad and less distinct, which makes it difficult to accurately determine the particle displacement and thus the velocity.
- **Large** *Δt*: Conversely, if *Δt* is too large, particles may travel beyond the boundaries of the interrogation window. This can lead to a loss of correlation because particles identified in the first frame are no longer present in the same window in the subsequent frame, resulting in inaccurate velocity measurements.

### Ensemble Correlation

Ensemble correlation is a technique in Particle Image Velocimetry (PIV) that improves the signal-to-noise ratio by averaging the correlation data across multiple frames or image pairs to enhance the accuracy of velocity field measurements (26,36). This is particularly important for complex or fluctuating flows with low “seeding” density as shown in Fig. 2. The time interval *Δt* between successive image pairs plays a significant role in determining the reliability of the correlation and, consequently, the precision of the flow velocity data. This is particularly useful in micro-PIV, where noise levels may be high due to the small scales of flow. The ensemble correlation function is typically expressed as (26,36):

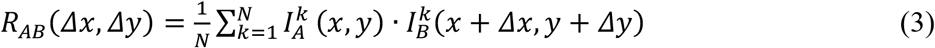

where,

- *R*_*AB*_(*Δx, Δy*): Ensemble correlation function between frames A and B for the displacement (*Δx, Δy*).
- *N*: Number of image pairs in the ensemble.
- 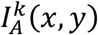: Intensity of the material elements in frame A at position (*x, y*) for the *k*^th^ image pair.
- 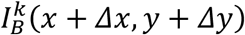: Intensity of the material elements in frame B shifted by (*Δx, Δy*) for the *k*^th^ image pair.

**Figure 2.**
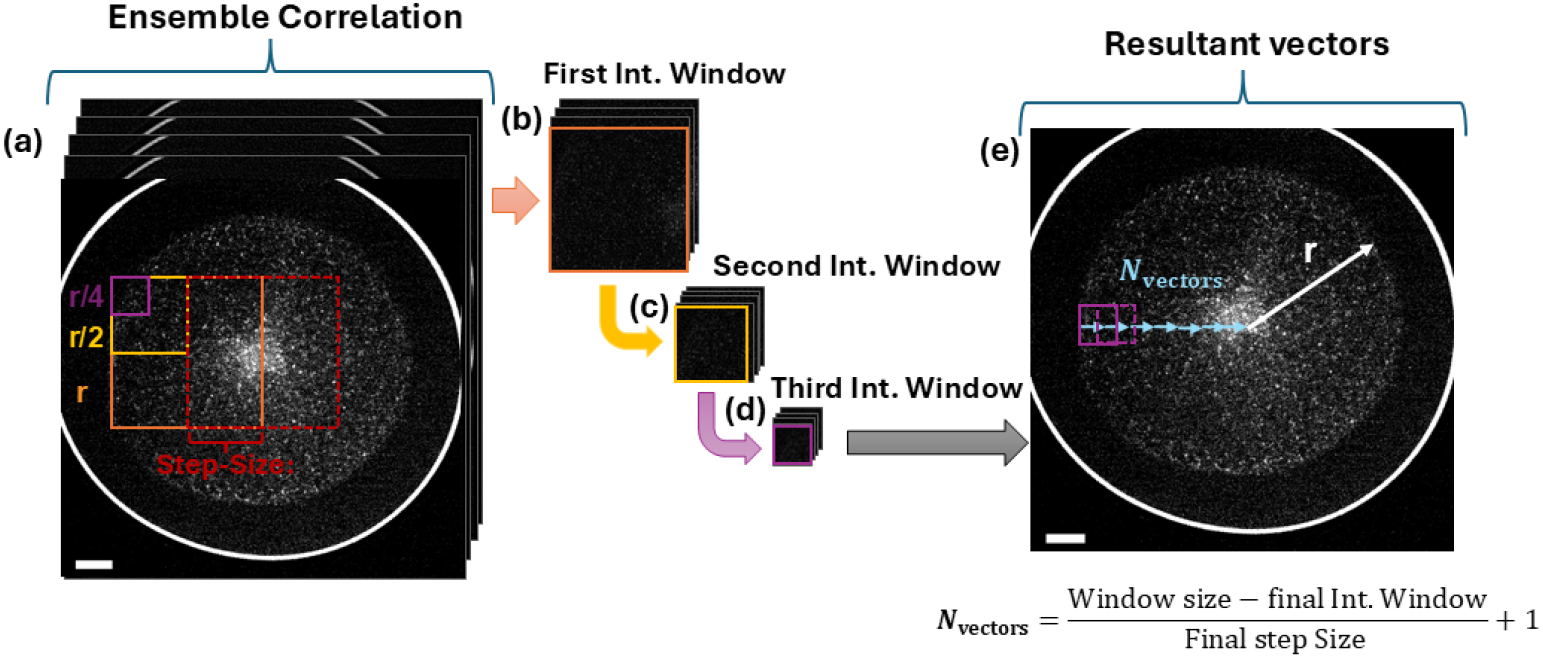
Ensemble correlation based Micro-PIV analysis of a contracting actomyosin gel (same gel as the one depicted in Fig. 1). (a) Ensemble correlation of multiple image pairs *N* (here 3) to average displacement (and velocity) measurements with three passes, each characterized by an interrogation window (Int. Window) of defined size: first (orange), second (yellow), and third (purple) with a step-size reduction of ½ (i.e., 50% shift or overlap) for every pass (here, shown only for first pass - dashed line red rectangle). (b) First interrogation window (*r* x *r* pixels, with radius *r* of the disc-shaped gel) used for initial displacement analysis; (c) Second interrogation window (*r*/2 x *r*/2) for further displacement calculation; (d) Third interrogation window (*r*/4 x *r*/4) for final displacement calculation; (e) Resultant displacement vector field obtained from the ensemble correlation process using these Int. Windows’ and step size, consists of *N*_vectors_ = 7, where here the gel radius is the window size, and the final Int. Window and step size are *r*/4 and *r*/8, respectively.

We note the following important steps:

1. **Cross-Correlation**: This expression computes the spatial correlation between two sequential images A and B, shifted by a vector (*Δx, Δy*).
2. **Ensemble Averaging**: Summing the correlations over *N* image pairs, reduces random noise and enhances the detection of true particle displacements.
3. **Peak Detection**: The peak of *R*_*AB*_ indicates the most likely displacement vector, which corresponds to the local material velocity.

During the first interrogation pass, larger interrogation areas are employed to capture the initial flow field, providing a coarse but stable estimate of displacement vectors, as seen in Figure 2. Subsequent passes, such as the second and third iterations, progressively refine the analysis by reducing the size of the interrogation window and step size. This iterative refinement enhances spatial resolution and vector accuracy by capturing finer flow details and correcting for any residual inaccuracies from earlier passes. Additionally, as the number of correlation frames increases, the averaging process improves statistical reliability, reducing noise and further refining the flow field. By optimizing both the interrogation window size (or area) and the number of passes, a balance is achieved between vector density (*N*_vectors_, denoting the number of vectors per gel radius) and resolution, enabling a comprehensive and accurate characterization of the flow dynamics.

Like for pair-correlation in conventional PIV, the choice of *Δt* between frames to be correlated is crucial for accurate velocity estimates. In addition, selecting the number of image pairs thoughtfully is essential to ensure accurate and reliable velocity measurements. Specifically, when *Δt* is small, the temporal resolution of the measurements is high, which is beneficial for capturing rapid changes in the flow. However, small values of *Δt* lead to small particle displacements, making it challenging to distinguish between true flow-induced motion and random noise. This can result in issues like peak-locking (sometimes called pixel locking (28)), where the displacements are insufficient to produce a meaningful correlation signal, ultimately affecting the accuracy of the velocity estimation. In contrast, increasing *Δt* allows for larger particle displacements, improving the ability to correlate particles over successive frames. However, if the displacement becomes too large, it may exceed the resolution capabilities of the imaging system, causing the correlation to degrade and leading to less accurate velocity estimates.

The number of image pairs *N* is another crucial parameter as it directly impacts the statistical reliability and accuracy of velocity measurements. Specifically, increasing the number of image pairs reduces random errors, enhances measurement accuracy, and boosts confidence in the computed velocity fields. When selecting this number, it is crucial to consider the intrinsic flow variability: flows exhibiting significant temporal fluctuations or turbulence necessitate more image pairs (also usually combined with smaller *Δt*) to adequately represent their statistical characteristics and achieve reliable measurements. Slower and laminar flows necessitate less image pairs and larger *Δt* might be sufficient to measure the more gradual changes in the flow.

### Image Pre-processing

To ensure the quality and reliability of data in Micro-PIV studies, the captured raw images undergo several enhancement steps. These steps are done in ImageJ Software.

First, background subtraction is performed to eliminate static noise and non-moving elements, such as reflections or debris, from the images. This process isolates the motion of material elements, thereby improving the clarity of their trajectories.

If the imaging system or experimental setup is affected by vibrations or unintended movements, stabilization techniques are applied to align consecutive frames, ensuring that the detected motion is attributed solely to fluid flow.

Additionally, in cases where the images exhibit blurriness due to optical distortions or scattering, deconvolution techniques can be utilized. Deconvolution enhances image sharpness and resolution by compensating for distortions introduced during the imaging process. Together, these steps, background subtraction, stabilization, and deconvolution as shown in Fig. 1(b) and described in the SI Supp. Note 1, play a crucial role in optimizing image quality and ensuring the accuracy of subsequent velocity and strain analyses in Micro-PIV experiments.

### Masking

In Micro Particle Image Velocimetry (Micro-PIV), masking is a crucial technique employed to isolate and analyze specific regions of interest within captured images as shown in Fig. 1(c). In that figure, the region of interest is the actomyosin gel disc. This method is especially useful for studying intricate flow dynamics or when focusing on specific areas within a microfluidic system. The masking process involves defining selected regions in the image for detailed analysis while excluding extraneous areas that could introduce noise or distort the results. Typically, this is accomplished using image processing, where regions of interest are delineated either manually or automatically, ensuring a more accurate and focused analysis of the fluid flow.

There are several reasons why masking is essential in Micro-PIV:

1. Focus on Relevant Regions: Microfluidic systems often contain intricate geometries or flow patterns. By masking out irrelevant areas, researchers can concentrate their analysis on the regions where the flow behavior is of particular interest. This allows for a more focused and efficient interpretation of the experimental data.
2. Noise Reduction: In some cases, certain parts of the image may contain artifacts or background noise that can affect the accuracy of the velocity measurements. By applying masks, these unwanted elements can be excluded from the analysis, leading to cleaner and more reliable results.
3. Improved Measurement Accuracy: Masking allows for precise control over the areas where velocity measurements are performed. By excluding regions with disturbances or low particle concentration, researchers can ensure that the analysis is conducted in areas where the flow is representative of the intended conditions. This enhances the accuracy and validity of the obtained velocity data.
4. Enhanced Computational Efficiency: By limiting the analysis to specific regions of interest, masking can reduce the computational burden associated with processing large image datasets. This can lead to faster analysis times and more efficient utilization of computational resources.

After pre-processing the images, we use MATLAB code to carry out the PIV analysis (36). The first step is masking the gel to avoid unwanted contribution from the outer background region. The procedure to carry this out along with Matlab code snippets are given in SI Supp. Note 2. The second step is to submit the masked images to the main PIV file, whose functionality is described in SI Supp. Note 3. For each gel we also evaluate the lateral contraction velocity *v*(*t*) by measuring the changes in gel radius *r*(*t*) with time.

## Results

We now present our key results from analyzing the spatio-temporal contraction dynamics of poroelastic actomyosin gels using the micro-PIV methods described above.

### Contraction dynamics follows characteristic poroelastic time scale

Figure 3 depicts the characteristic time-dependent dynamics of the contractile poroelastic actomyosin gel depicted in Figs. 1 and 2. The reduction in gel disc radius *r* as a function of time indicates a rapid initial decrease followed by a slower decay phase (main panel). The corresponding gel edge radial contraction velocity *v*(*t*) exhibits an initial rapid contraction, characterized by a linear increase in contraction velocity up to maximal value (*v*_*max*_). This initial linear phase is followed by an exponential decay phase, *v*(*t*) = *v*_*max*_*e*^−*t*/τ^, where *τ* is the characteristic relaxation time (13). The decay constant is proportional to an effective friction constant that accounts for the permeation of water through the actin gel pores and inversely proportional to the gel elastic modulus. At the end of this phase the gel reaches a mechanically stable state where the active contractile stresses exerted by myosin motors in the gel are balanced by the gel elasticity.

**Figure 3.**
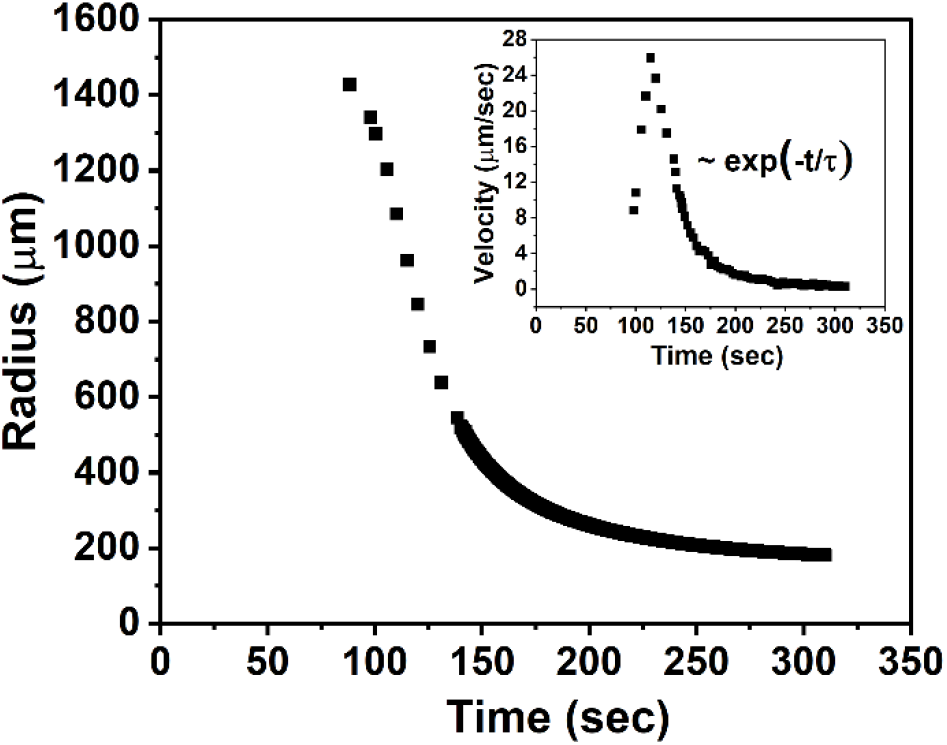
Time-dependent evolution of actomyosin gel dynamics of the gel disc shown in Fig. 1. The main plot shows the reduction in gel radius *r* versus time from its initial value (*R* = 1600μm). The inset presents the corresponding gel edge radial contraction speed *v*(*t*) characterized by an initial linear growth phase, followed by an exponential decay phase with speed *v*(*t*) = *v*_*max*_*e*^−*t/τ*^, *τ* = 28s and *v*_*max*_ = 26μm/s. Images are acquired with a temporal resolution of 0.1s.

We employed Micro-PIV to analyze the contraction dynamics and velocity field of the actomyosin gel. Initially, we investigated how varying the time interval *Δt* affects the displacement field while maintaining the number of frame pairs constant (*N* = 299), as depicted in Figure 4. Here, *Δt* was varied between 0.1 to 0.5s, which corresponds to 1 to 5 frames between consecutive images. At *Δt* = 0.1s, the displacement vectors are small but that still captures the overall radial flow of the contracting gel. However, increasing *Δt* to 0.3 and 0.5 s produces larger displacement vectors, which, while expected, also lead to challenges in maintaining correlation accuracy, particularly at the highest *Δt*. Additionally, for *Δt* values of 0.3 and 0.5 s, the overall ensemble correlation time (defined as *N* × *Δt*) is 90 and 150 s, respectively – 3 and 5 times longer than the characteristic time scale (∼30 s) governing the contraction dynamics in both the linear and exponential regimes (Fig. 3). This suggests that for accurate measurement of displacement vectors, it is essential to adjust not only *Δt* but also the total ensemble correlation time.

**Figure 4.**
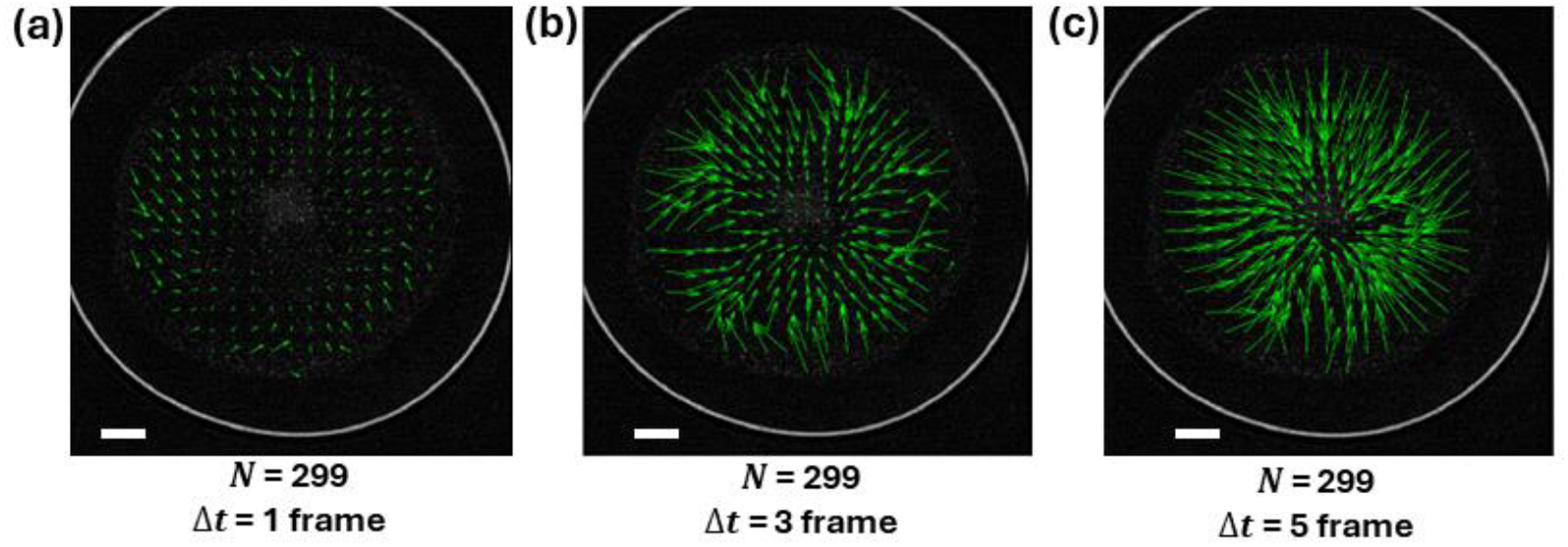
Effect of varying *Δt* (in number of frames) on the displacement vector field resolution with an ensemble of 300 correlation frames (i.e., *N* = 299 image pairs). Images are acquired with a temporal resolution of 0.1s per frame. (a) At *Δt* = 1, the displacement vectors are minimal, leading to a low magnitude field. (b) At *Δt* = 3, the displacement vectors are more pronounced, capturing moderate flow dynamics. (c) At *Δt* = 5, the displacement vectors are further amplified, showing higher displacement magnitudes and enhanced flow structure. The increasing *Δt* reveals clearer vector distributions but may introduce challenges in maintaining correlation accuracy for larger displacements. Analysis is performed on the gel depicted in Fig. 1. Bars are 300μm.

### Adapting the interrogation window sizes to the size of the mask – effects of number of passes and step size on displacement vector field

The sizes of the interrogation windows are systematically adjusted to match the gel’s dimensions at each time point, ensuring precise analysis within the defined region of interest. In the first interrogation pass, large windows capture the overall flow-field and provide a coarse but robust estimate of displacement (Fig. 2). Subsequent passes, each with progressively smaller window and step sizes, refine these estimates. This adaptation directly affects both the spatial resolution and the density of the resulting displacement vectors (Fig. 5).

**Figure 5.**
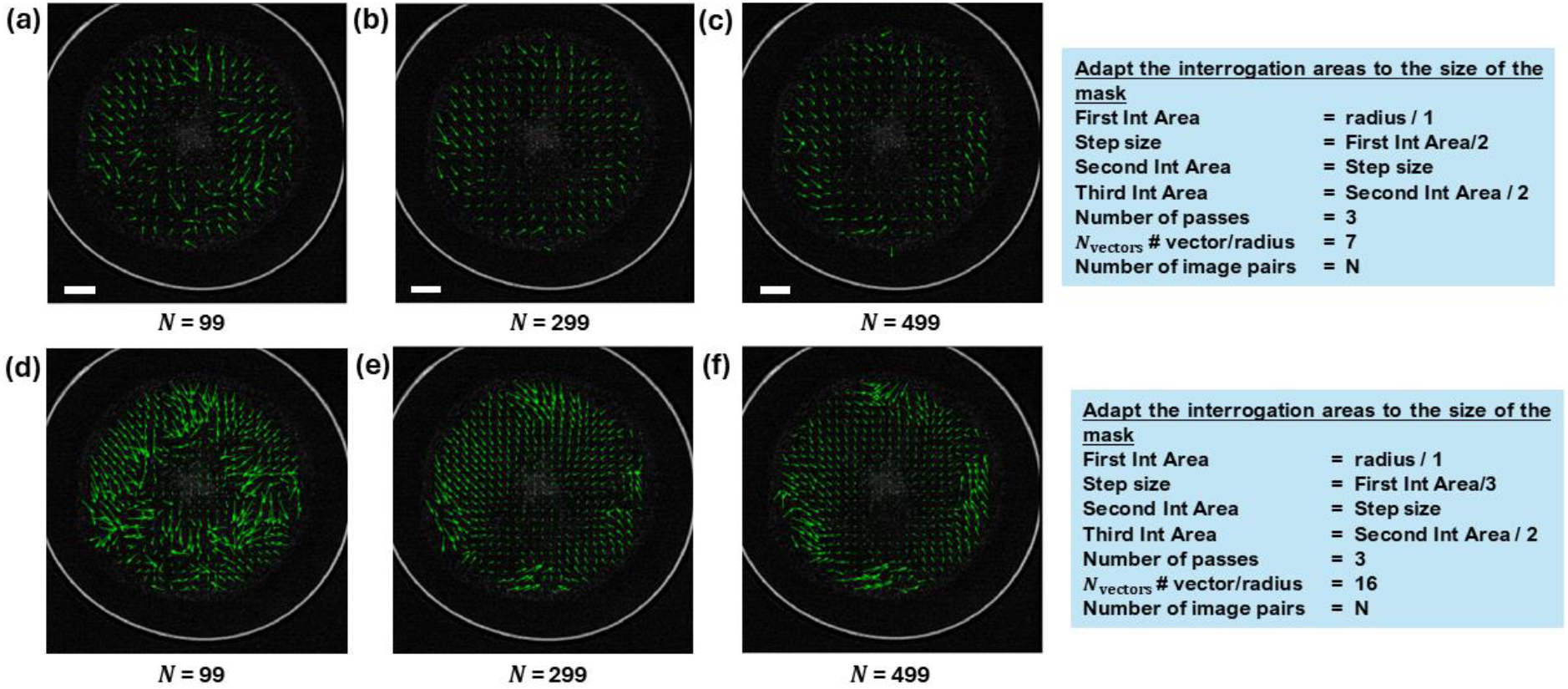
Effects of interrogation window area (size), step size, and number of image pairs on displacement vector field. Displacement vector fields obtained through ensemble correlation with varying frame pairs and interrogation window sizes and step sizes. Analysis is performed on the gel depicted in Fig. 1. (a–c) Correlation frames of 100, 300, and 500 respectively, which corresponds to number of image pairs of 99, 299, and 499 respectively, with step size scaling of Interrogation size/2. (d–f) Same correlation frames as in (a-c), with step size scaling at Interrogation size/3. The number of passes for all cases is fixed at 3. The First interrogation area is adapted to the mask size (i.e., radius). The time interval is fixed at *Δt* = 0.1s.

We analyze the contraction dynamics of the gel in Fig. 3 by applying 3 interrogation passes. The initial window size is chosen to match the gel’s radius (set by the mask) at each time point (Fig. 1c). Figure 5 illustrates how varying window size, step size, and the number of image pairs (*N*) used in ensemble correlation (of *Δt* = 0.1s) impacts the computed displacement field. With a larger step size, displacement vectors are larger but less densely distributed yet still captures the overall radial flow of the contracting gel when using 99 image pairs (Fig. 5a–c). Increasing *N* from 99 to 299 improves correlation accuracy; further increase in *N* to 499 yields negligible improvement. Conversely, reducing step and window sizes, as expected, increases the density of resulting displacement vectors *N*_vectors_ from 7 to 16, but lowers accuracy, likely because smaller displacement vectors. This effect is most significant at low *N*. Although higher *N* partially mitigates this inaccuracy, it remains suboptimal even at the largest sample size. Therefore, balancing interrogation window size, number of passes, and image-pair count is essential to optimize both vector density and measurement accuracy for comprehensive characterization of the gel’s flow dynamics.

In the context of PIV or similar image analysis techniques, adapting the interrogation windows ensures that the analysis can accurately capture the flow or motion within the masked region. This part of the script to adapting the interrogation areas (windows) to match the size of the mask and the procedure to carry this out along with Matlab code snippets are given in SI Supp. Note 4. The purpose of this code is to define a series of interrogation areas of progressively smaller sizes. These sizes are related to the radius of the mask, ensuring that the interrogation windows are appropriately scaled for the analysis of the masked region. Smaller interrogation windows can capture finer details of the flow or motion but can be noisier while larger windows can provide an overview. In practical terms, setting interrogation areas in this way allows the PIV algorithm to:

- Start with a larger area to capture broad motion or flow patterns.
- Progressively use smaller areas to refine the analysis and capture finer details.

This multi-scale approach can improve the accuracy and resolution of the PIV results. By progressively reducing the size of the interrogation areas, the script can adapt to the size of the mask and improve the detail and accuracy of the analysis. These parameters are for the interrogation window and must change as per the gel or how many vectors you want in the PIV output.

### Displacement and strain analysis between frames

The actomyosin gel is a poroelastic material comprising an elastic network permeated with solvent. The timescale over which the gel displacement vectors are measured in PIV (∼0.1s) is much shorter than the poroelastic stress relaxation time scale given by the timescale of contraction of the gel disk (see Fig. 3). The gel deformation between two frames can thus be described by the linearized strain rate tensor, constructed from the displacements measured in PIV (37) The displacement vectors obtained from the PIV are located at regularly spaced lattice points (Fig. 6a), from which the gel strain can be calculated using finite differences.

**Figure 6.**
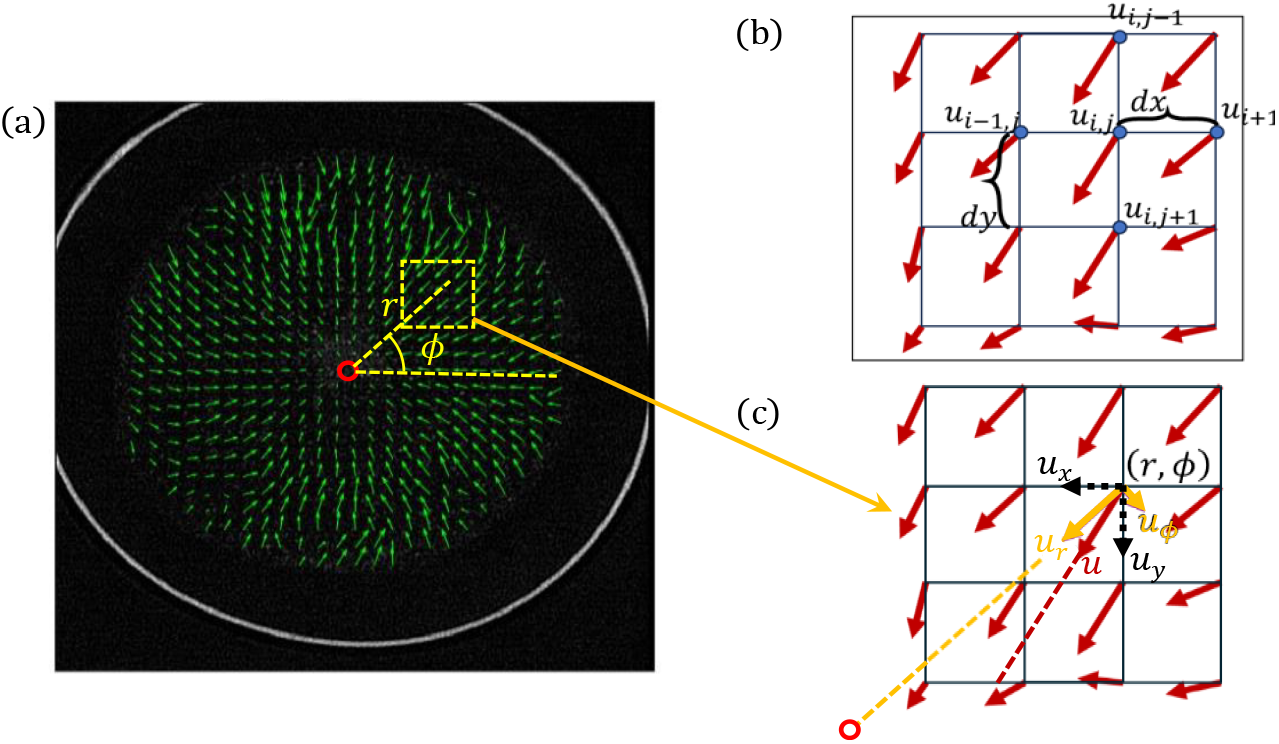
(a) Representative view of the gel disk shown in Fig. 1, with superposed lattice of displacement vectors obtained from PIV and choice of coordinate system. (b) The cartesian components of the strain tensor are obtained by taking derivatives of the displacement using the central difference method on the square lattice. (c) The displacement vectors at the lattice points, measured in cartesian coordinates, can be converted into corresponding radial and azimuthal components, *u*_*r*_ and *u*_*ϕ*_. (a and c) The origin of the coordinate system is chosen to be at the geometric center of the gel (red circle), after subtracting the rigid body displacement of the center of mass if required.

The strain tensor is the gradient of the displacement vectors and encodes geometric information on the different deformation modes of local slices of the gel: stretching or compression, shear, and rotation (38). We note that this is more commonly computed as a strain rate in usual PIV analysis, but we here do not divide by the time interval, *Δt*. This gives the information of material deformation over a small-time interval.

### Calculation of local strain

- The displacement vectors are obtained for each lattice point (i, j) from PIV in cartesian form, i.e. *u*_*x*_ and *u*_*y*_ (Fig. 6b).
- To obtain the different components of the strain tensor, i.e. *ε*_*xx*_, *ε*_*xy*_, *ε*_*yx*_ and *ε*_*yy*_ at location (i, j), we consider the displacements *u*_*i*−1,*j*_, *u*_*i,j*−1_, *u*_*i*+1,*j*_ and *u*_*i,j*+1_ i.e. displacements at locations (i − 1, j), (i, j − 1), (i + 1, j) and (i, j + 1) respectively as shown in Fig. 6b.
- Thereafter, we use the three-point central difference method, to obtain the strain tensor components.

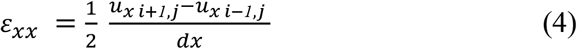

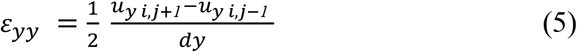

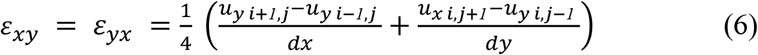

### Estimation of local displacements and strains in polar coordinate basis

Most of the gels are circular disks contracting in an apparent axially symmetric manner. Hence, we represent the strain in 2D cylindrical polar coordinates to better address this geometric feature of the gels. The steps for converting the cartesian strain components to azimuthal and radial components are outlined below.

- Locating the centroid of the gel: The masked lattice points are used to locate the centroid (or geometric center) of the gel. The *x* and *y* coordinates of the centroid are given by the arithmetic mean of the coordinates of all the individual lattice points in the masked region of the gel (Fig. 1c, circular black region).
- The center of the gel is set as the origin, i.e. `(0,0)` as shown in Fig. 6 (a) and polar coordinates of the lattice points are calculated with respect to this origin. The distance between the centroid and each lattice point is given by separation length (r). The angle *ϕ* is made by the position vector of each lattice point from the centroid with respect to the horizontal:
- Displacements in polar form are calculated
- The strain is calculated in polar coordinates using the equations below

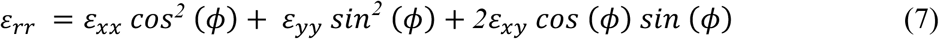

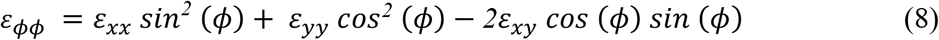

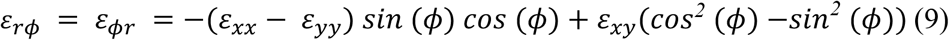

### Bulk or Area strain maps as a measure of local contraction

The representative 2D heatmap of bulk or area strain (shown in Fig. 7) is obtained by tracing filled contour plots of the trace of the strain tensor (*ε*_*xx*_ + *ε*_*yy*_) at each lattice point. This measures the net area deformation (contraction or expansion) at a given location in the gel. Positive (negative) values correspond to expanding (contracting) regions. In Fig. 7, to visualize the sign of strain, each individual positive or negative area strain value is normalized by the maximum positive and negative values in the whole gel at that specific time. The MATLAB code used to generate the 2D heatmap of bulk area strain in Figure 7 is described in SI Supp. Note 5.

**Figure 7.**
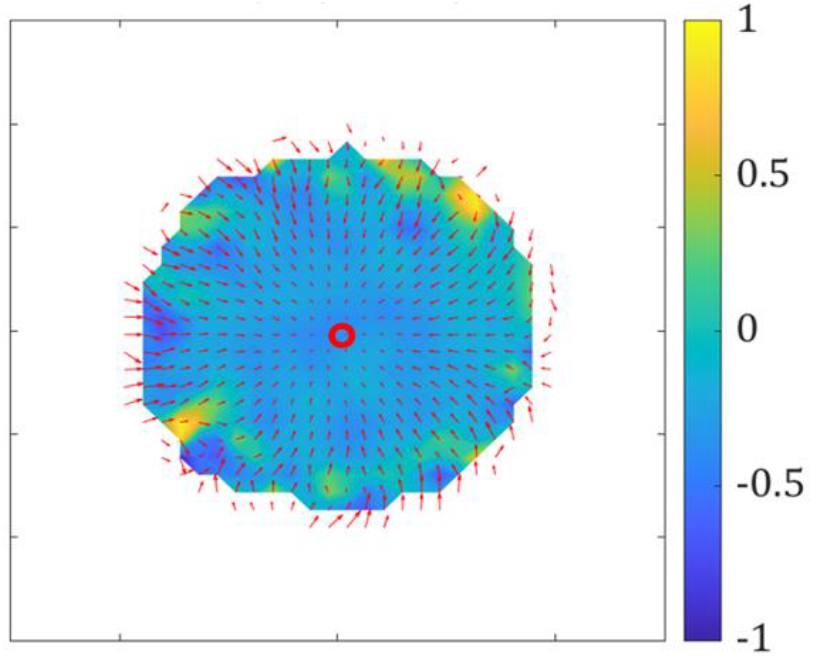
Area strain (trace of the strain matrix) of the gel shown in Fig. 1. The trace of the strain between two frames is shown as a colormap (blue: negative or contraction, yellow: positive or expansion). The red arrows show the displacements obtained from PIV for the chosen Δ*t*. The inner region of the gel undergoes rapid contraction, as indicated by dark blue regions. Closer to the boundary, we find patches of stretched regions shown by the yellow hotspots (The strains are normalized by the maximum positive and maximum negative values measured in the entire gel, respectively, at that specific time).

Here the centroid of the gel is demarcated by the red circle, the PIV arrows are shown in red, and the area strain is shown as a heatmap. The contracting regions, i.e. regions with negative strain, are shown in blue, while the stretched regions with positive strain are in yellow. We see that the inner region is mostly blue corresponding to contraction, while there are patches of stretched regions (yellow) near the boundary of the gel. This indicates non-uniform or anisotropic contraction of the gel, where the boundary contracts at a different rate and direction than the interior. While a representative frame from the PIV analysis is shown here, the time course of evolution of the area strain for this gel is shown SI Fig. S1, using additional selected frames.

We may follow a similar procedure to analyze 2D heatmaps of other components of the strain. The radial strain *ε*_*rr*_, and azimuthal strain *ε*_*ϕϕ*_, reveal the stretch or compression in radial and azimuthal directions respectively. For perfectly isotropic contraction, they are equal. The difference between the radial and the azimuthal strain components reveals the extent of anisotropy in the motor-generated active stresses.

### Principal strain component analysis

While the trace of the strain gives just the magnitude of expansion or contraction, an analysis of the eigenvectors of the strain tensor reveals the *principal directions* of the expansion or contraction. The eigenvalues of the strain tensor are given by,

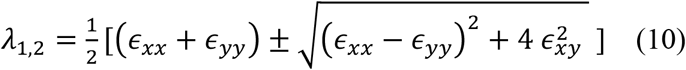

with corresponding eigenvectors:

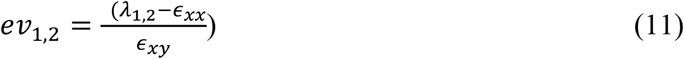

As shown in the schematic in Figure 8, the strain in general contains stretching/compression but also deviatoric or shear components that correspond to shape change. Transforming to the eigen basis removes the simple shear component and gives orthogonal directions corresponding to the maximum and minimum stretching/compression, respectively. In general, these are not necessarily the radial and azimuthal directions.

**Figure 8.**
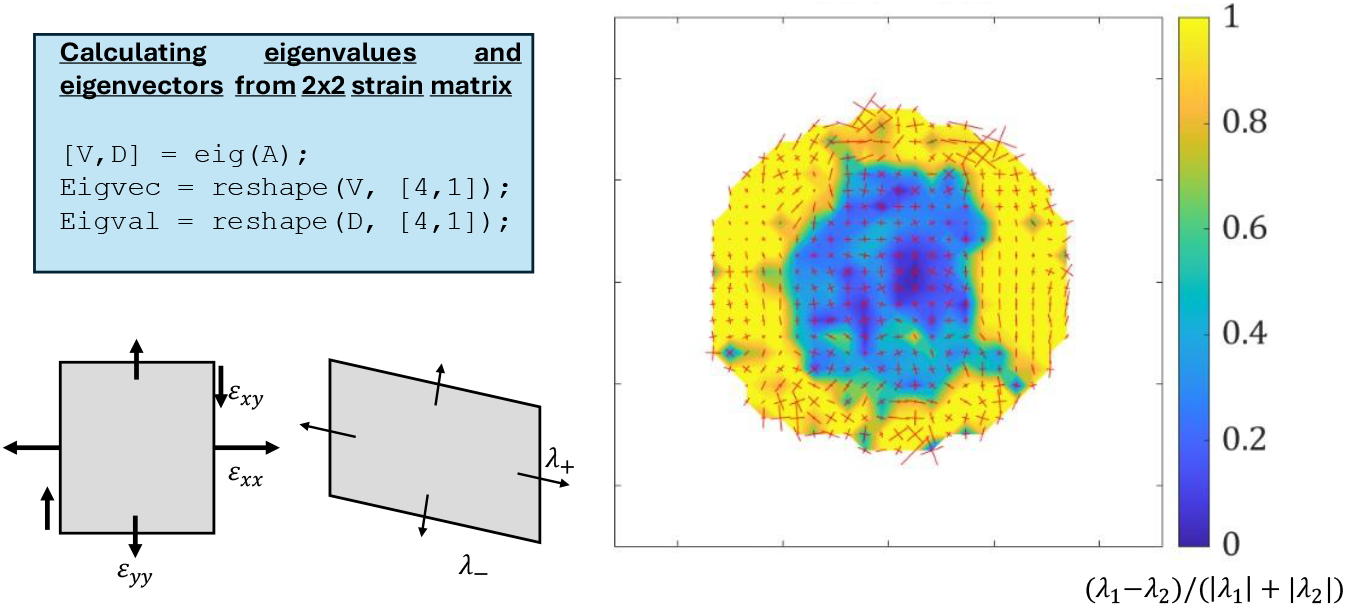
Eigenvalue and eigenvector analysis of the strain tensor gives the principal directions of contraction and stretching of the gel depicted in Fig. 1. (*Left)* Schematic illustrating the computational procedure and intuitive meaning of principal strain components (maximal stretch/compression directions). (*Right*) The color map shows the normalized difference of the two strain eigenvalues for a representative frame from PIV of a circular gel. The vector map shows the two orthogonal eigenvectors scaled by the magnitude of its corresponding eigenvalue. The plot indicates an isotropic inner region (blue) where both eigenvalues are almost equal, and an anisotropic outer region (yellow). The outer region is characterized by increased alignment of strain along the azimuthal direction, with some local instances of high radial stretching.

In Figure 8, we show the normalized difference of the two eigenvalues, (*λ*_1_ − *λ*_2_)/(|*λ*_1_| + |*λ*_2_|) as the color map. A lower value (closer to 0) suggests a high degree of isotropy, while a bigger value (closer to 1) indicates that one eigenvalue is dominant, corresponding to significant anisotropy. We also show both orthogonal eigenvectors in the vector plot, scaled by their respective eigenvalues. We observe no apparent alignment of strain in the inner region of the gel (blue), where both eigenvalues are nearly equal. This corresponds to isotropic contraction in the inner region and is consistent with the overall negative area strain seen in Figure 7. Close to the boundary, the alignment of eigenvectors is predominantly azimuthal, with one eigenvalue significantly higher in magnitude than the other. This suggests azimuthal alignment of actin bundles and therefore, of contractile actomyosin force dipoles, at the gel boundary. There are a few instances of radially aligned eigenvectors accompanied by large positive eigenvalue at the boundary. This suggests radial stretching of the material in response to azimuthal contraction imposed by actomyosin forces, because of the Poisson effect (37) .This effect provides evidence for the elastic response of the gel to active stresses, at short timescales.

While Figure 8 shows a representative frame from the PIV analysis, the SI Fig. S2 shows similar plots for a sequence of frames. Overall, we observe azimuthally aligned eigenvectors close to the boundary of the gel (yellow) that suggest azimuthal contraction, while the inner region (blue) acts as an isotropic core of contraction.

### Radial and azimuthal displacement and strain obtained from axisymmetric averaging

The gel in Figure 1 which we analyze here appears to contract towards the centroid with all displacement vectors oriented radially. Hence, we expect that the local azimuthal displacements to be very small and assume that the gel shows axisymmetric contraction. We consider circular annuli of equal thickness, *δr* from the gel boundary to the centre (a total of 25 annular bins) (Fig. 9a). The bin width *δr* is calculated at every frame and is given by 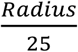 (from Fig. 3). In each of these annular bins, there are multiple PIV lattice points. We calculate the mean and standard deviation of the radial and azimuthal displacements in each of these annular bins. To validate our assumption of axisymmetric contraction, we consider azimuthal displacement *u*_*ϕ*_(*r*) as a function of the normalized distance from the centre of the gel (Fig. 9b). It fluctuates closely to 0, suggesting no apparent average azimuthal displacement, which suggests axisymmetric contraction. For angle-averaged radial displacements in annular bins, we obtain an inner region which shows linear negative dependence on distance from the center. Closer to the boundary, the radial displacement profile curves up to form a “hockey stick”-like shape suggesting radial stretching. This stretching close to the boundary correlates with the observed positive patches of area strain in Figure 8.

**Figure 9.**
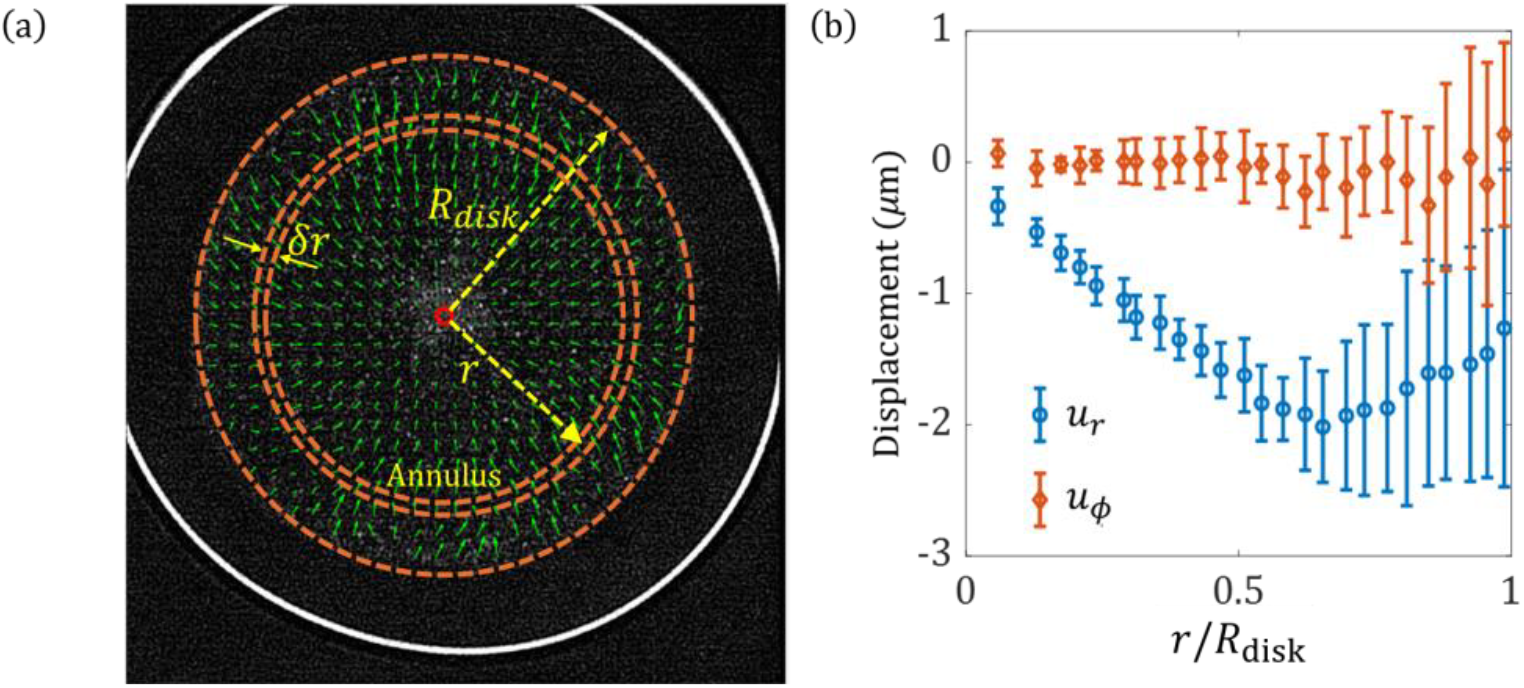
(a) Circular annular bins are constructed around the centroid of the gel disc shown in Fig. 1. The radial and azimuthal components of the displacements from Fig. 6c are averaged over all angles in each circular bin. The bin width, *δr* depends on the gel disk radius at that frame, 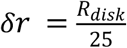. In other words, there are 25 circular annuli from center to the boundary. (b) The angle-averaged azimuthal displacement (*u*_*ϕ*_) does not change with distance from the center of the gel and fluctuates close to 0. The angle averaged radial displacement (*u*_*r*_) increases linearly in magnitude from the center in the inner region and decreases closer to the boundary in the outer region.

### Analyzing gels with non-axisymmetric displacement field

- Some gels exhibit rigid body motion, that is displacement of their center of mass, as they contract. The displacement of the gel is no longer axisymmetric about its center of mass which we can verify by constructing concentric annuli around this point. We see for example in Figure 10 that some of the displacement vectors in these annuli are not oriented radially inward. We can instead identify an apparent centre of displacement (*CoD*) for such gels from the displacement vectors and estimate the radial profiles of displacements and strains with respect to this special point (*CoD*). We note that all deformation of the gel is encoded in the strains, which are gradients of the displacement. The displacement field, by itself, does not provide any fundamental insights into gel contraction. Nonetheless, by subtracting the rigid body or center of mass (*CoM*) motion, and subsequent averaging, we show that we recover universal trends in the radial gel displacement profiles. To determine the location of the centre of displacement (*CoD*), we first calculate the mean displacement of the gel, i.e. the average of all the PIV displacement vectors

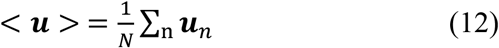 This gives the displacement of the center of mass, and thus the rigid body translational motion of the entire gel. When the *CoD* of the gel is not at its centroid, there is a rigid body motion associated with the gel. By subtracting the rigid body motion of the gel, the PIV displacement is transformed although the strain components are expected to remain the same.
- We now outline the procedure to locate the center of displacement relative to the center of mass. The general idea is to find the point where the displacement vector of a given lattice point intersects that of the *CoM* and then average over all such lattice points. Like all lattice points in the gel, the *CoM* also moves towards the *CoD*, i.e.

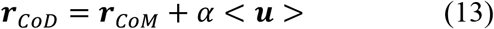

where *α* is an unknown constant, and the above relation states that the *CoM* displacement is parallel to the separation vector connecting the *CoM* to the *CoD*. This is illustrated in the schematic in Fig. 10a. To determine the unknown *α*, we consider the intersection of the mean displacement vector and all displacement vectors in the gel. For a general lattice position within the gel, whose position relative to the *CoM* is given by **r**, the corresponding displacement vector is also assumed to point towards the *CoD* (see Fig. 10a), giving

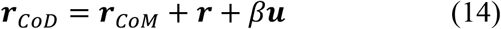

Combining the above two relations in Eqs. 13 and 14, we get: *α* < ***u*** > = ***r*** + *β****u***. To eliminate the second unknown *β*, we take the cross product of both sides of this equation with the displacement vector **u** to get,

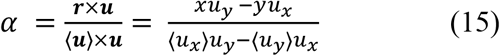

where on the right-hand side, we have explicitly written the expression for *α* in terms of *x* and *y*, the cartesian coordinates for the position of a lattice point relative to the *CoM*, and its associated displacement components, *u*_*x*_ and *u*_*y*_. Due to spatial fluctuations in the PIV data, we obtain a distribution of values for *α* shown in Fig. 10b. We drop the extreme outliers which arise from oppositely directed vectors, and are thus unphysical, and consider instead the sharp peak (median) value, which gives the *CoD*. Figure 10 shows the result of this calculation and demonstrates how the centroid (*CoM*) of the gel is different from the center of displacement (*CoD*).
- In Fig. 10c, we plot the displacement vectors after subtracting the *CoM* displacement, i.e. the rigid body motion, *u* − ⟨*u*⟩.
- We now transform to polar coordinates relative to this calculated center of displacement as the new origin. The radial direction for each displacement vector in the lattice is now relative to the *CoD*. We calculate profiles of the radial displacement *u*_*r*_ by averaging over annular bins as described before. We compare in Figs. 10d and 10e these radial displacement profiles centered on the *CoM* and *CoD* respectively. As expected, the displacement profiles are less noisy, and the characteristic trends much more clearly seen in Fig. 10e, where we used the *CoD* as our point of reference.
- The method of locating the center of displacement is also important for irregular gels that deviate from a full circular contour. This includes for example a gel whose boundary forms part of a circular arc (Fig. 11) and for gels that do not have a well-defined centroid. We can determine the radial profiles of displacements and strains in this circular sector enclosed by the arc (Fig. 11b,c). Further analysis of the trace of the strain and the principal components of the strain at different frames for the gel shown in Fig. 11, are provided in the SI Figs. S3 and S4, respectively. Fig. S4 shows evidence for strong anisotropy in the strain field. In almost all cases seen, the dominant strain eigenvector is azimuthal, providing strong evidence for azimuthal alignment of the force-producing actomyosin fibers near the boundary of the gel.

**Figure 10.**
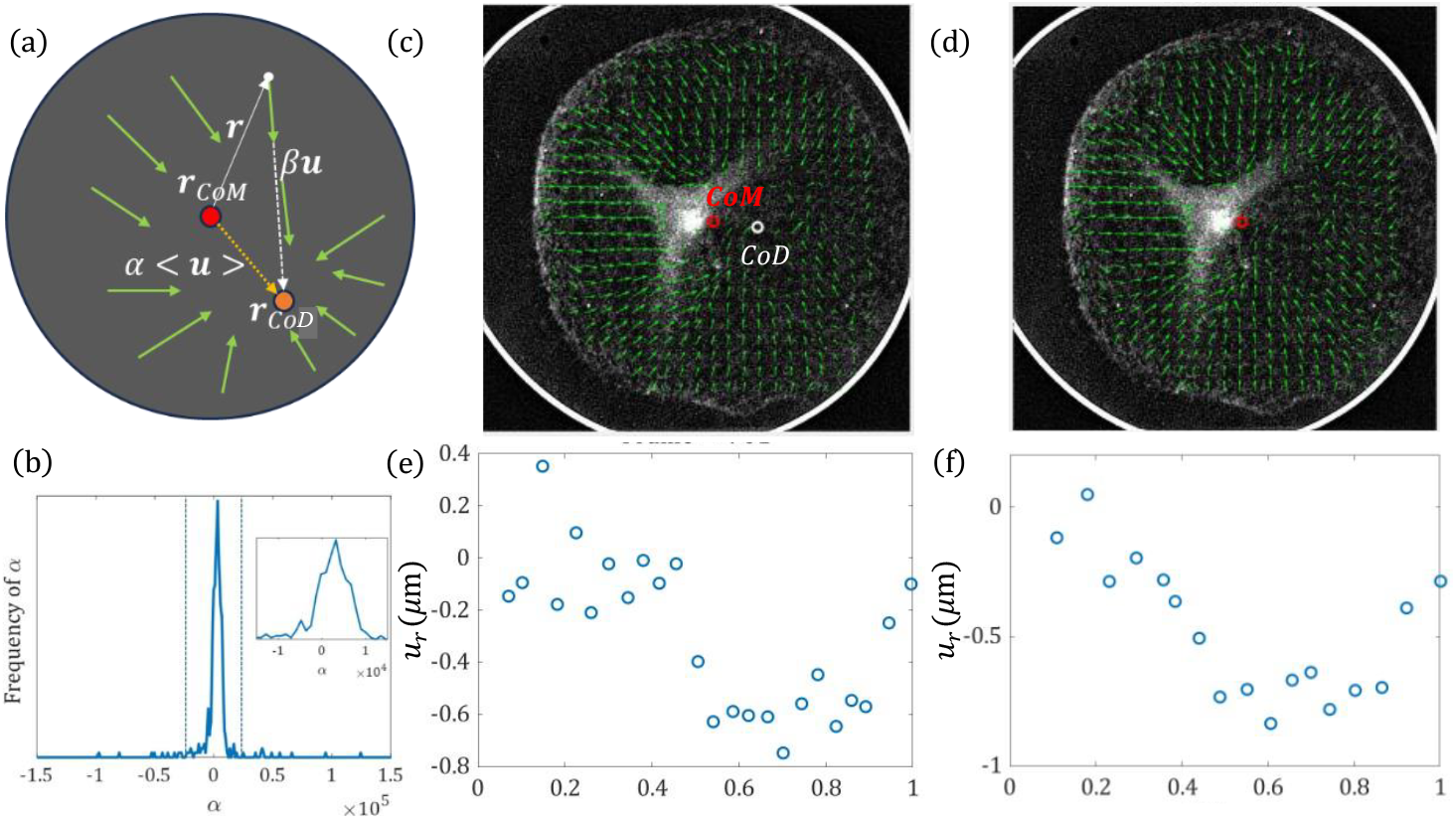
(a) When the center of mass (*r*_*CoM*_) of the gel displaces, the displacement vectors apparently converge to a shifted center of displacement (*r*_*CoD*_). (b) The distribution of the shift factor α (Eq. 15) over different lattice vectors from the PIV, with the inset focusing on the strong peak. (c) A sample PIV with centroid of the gel shown by a red circle while the apparent center of displacement to which the displacement vectors converge is shown by a white circle. The latter is calculated from the median intersection point of the PIV vectors (see Eq. 14 in main text). (d) The PIV map is transformed after subtracting the displacement of the center of mass which represents the rigid body motion of the gel. The center of displacement now coincides with the centroid shown by the red circle. (e) A very noisy radial profile of the radial displacement is obtained by binning with respect to the centroid of the gel, since some of the displacement vectors are oriented away from the centroid close to the center of the gel. (f) Noise is reduced when the radial displacement profile is calculated relative to the center of displacement. This is because in this transformed coordinate system, the PIV displacement vectors point towards the *CoD* restoring the axisymmetry of the displacement field. The radial binning procedure now works as expected. This restores the characteristic radial displacement profiles observed in the more symmetrically contracting gels, suggesting the same underlying physical mechanism in the gel contraction.

**Figure 11.**
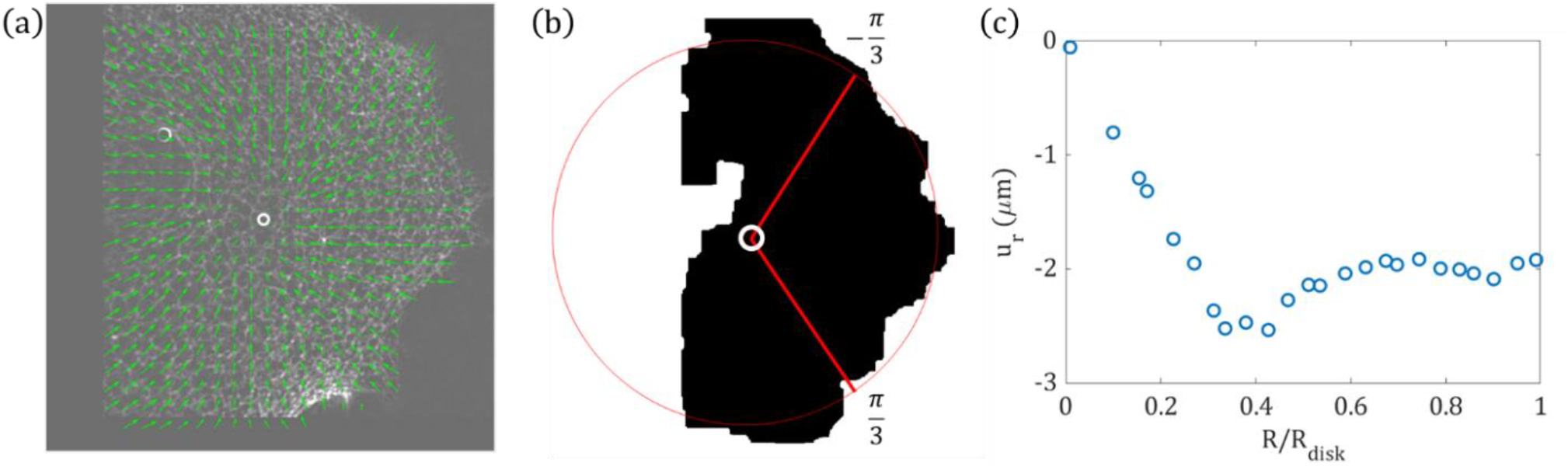
(a) PIV map of an irregular shaped gel with displacement vectors pointing to the center of displacement (white circle). (b) We consider a sector of the gel which forms a circular arc at the boundary between angles 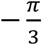 to 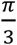 with respect to the center of displacement. The radial profile is estimated for the gel within the sector enclosed by the circular arc. (c) The radial profile of radial displacement for the gel within the circular sector is shown here.

From Figs. 10f and 11c, we see that gels that have moving geometric centers as well as of non-circular shape, give rise to qualitatively similar radial displacement and therefore, strain profiles.

## Conclusion

In conclusion, we establish and validate the methodology for adapting conventional micro-PIV to the dynamics of complex active gels. We showed how noise can be reduced in the imaging data both during pre- and post-processing the input and output of the micro-PIV analysis, respectively. Importantly, we showed how an optimal time interval can be chosen between the pairs of images to be correlated over to reliably calculate the velocity field. This choice relates to a physical timescale, in this case that of the poroelastic dynamics of gel contraction (13,15,35,39,40).

The analysis of strains from displacement data as well as averaged radial profiles of displacement reveal consistent features across gels of diverse initial and final shapes. This suggests that our method is robust, and uncovers common features in the dynamics, and therefore stress distribution, in these gels. Importantly, our work reveals that all gels contract isotopically in their inner region, but the principal strains are aligned azimuthally at the boundary. This correlates with the alignment of actin fibers observed at the boundary in fluorescence microscopy images (14). Further, we have shown that the analysis can be extended to gels whose contraction is apparently non-axisymmetric and that do not form a fully circular shape. We note that while PIV is a commonly used technique to identify local velocity and contraction rates in cytoskeletal networks (32,33,41–43), many of these prior works utilize traditional PIV methods where it was sufficient to correlate between just one image pair. On the other hand, we utilized correlation across an ensemble of image pairs to reliably extract global displacement field for our rapidly contracting gels. This method may thus in future be applicable to other cytoskeletal gels, including actomyosin-microtubule composites.

In summary, this research highlights the necessity of modified micro-PIV approaches for studying complex fluids, specifically poroelastic active materials with large, autonomous contraction. It thereby advances our understanding of active cytoskeletal dynamics and self-organization. The findings have broad implications for cell motility, morphogenesis, and active matter physics.

## Supporting information

Choudhary SuppInfo

## Acknowledgments

We thank Efi Efrati for useful discussions and for critical reading of the manuscript.

## Author contributions

SC Developed analytical methods for Micro-PIV analysis, prepared figures and movies, and wrote the manuscript. SB analyzed the Micro-PIV data, developed methods for strain analysis, prepared figures and movies. YA and DSS developed image analytical tools for data quantification. GL performed experiments and analyzed the experimental results. KD designed methods for strain analysis and wrote the manuscript. ABG Designed and developed the experimental system, developed analytical methods for data quantification, and wrote the manuscript.

## Data availability

All data are available in the manuscript or the supplementary materials.

