## Supplementary material for "Investigating active dynamics of contractile actomyosin gels with Micro Particle Image Velocimetry (Micro-PIV) analysis": Choudhary SuppInfo

**[3] Department of Physics, University of California, Merced, CA 95340 USA**

**[4] Department of Physics, SRM University-AP, Amaravati 522240, Andhra Pradesh, India**

*These two authors contributed equally to this work

#### Supplementary Note 1. Pre-processing images

**Step 1: Background Removal (in ImageJ Software)**

1. **Open the Entire Series as an Image**:
   - **Procedure**: Load the entire time-lapse series of images into the image analysis software (e.g., Fiji/ImageJ).
   - **Rationale**: This allows for batch processing of the entire image series. It also ensures that all frames are treated uniformly during preprocessing. Loading the entire image series into the analysis software enables uniform treatment and batch processing, ensuring consistency throughout the preprocessing steps.
2. **Subtract Background**:
   - **Procedure**: Navigate to `Image -> Process -> Subtract Background` in the software.
   - **Rationale**: Background subtraction is essential for removing fluorescence signal from actin monomers in the background, thereby enhancing the visibility of actomyosin structures. This process improves contrast, making these structures more distinguishable from the background, which facilitates more accurate downstream analysis by eliminating noise and focusing on the relevant features.

**Step 2: Stabilization (in ImageJ Software)**

1. **Install the StackReg Plugin**:
   - **Procedure**: Download and install StackReg Plugin.

- **Rationale**: The StackReg plugin is essential for aligning the frames of the image stack, correcting any drift that may have occurred during image acquisition. This ensures accurate alignment and consistency across all frames, which is crucial for precise analysis.

2. **Set the Reference Frame**:
   - **Procedure**: Open the background-subtracted time-lapse series. Move the time slider to the last frame to set it as the reference for stabilization.
   - **Rationale**: Selecting the last frame as the stabilization reference is crucial for achieving consistent alignment across all frames of the time-lapse series. By using a single reference point, any variations or shifts between frames due to minor movements or drift during image acquisition can be minimized. This approach ensures that all subsequent frames are aligned relative to this stable reference, thereby enhancing the overall stability and coherence of the time-lapse sequence. This stability is essential for accurate quantitative analysis of dynamic biological processes such as movement or changes in cellular structures, where precise spatial and temporal alignment is critical for meaningful interpretation and measurement.
3. **Apply StackReg Transformation**:
   - **Procedure**: In the software, go to `Plugins -> StackReg -> Transformation -> Translation`. Click `OK` to apply the transformation.
   - **Rationale**: All frames in the series are aligned with respect to the reference frame, effectively stabilizing the entire image stack. By minimizing frame-to-frame shifts caused by motion or drift during acquisition, the alignment ensures consistency and accuracy throughout the time-lapse sequence. This stabilized image stack is critical for reliable and precise subsequent analysis, enabling consistent measurement and comparison of dynamic processes over time.
4. **Save the Stabilized Image**:
   - **Procedure**: Save the stabilized image series as individual frames for better file handling.
   - **Rationale**: Saving the stabilized series ensures that the processed frames are preserved for further analysis or processing steps.

**Step 3: Deconvolution (in ImageJ Software)**

1. **Generate a Point Spread Function (PSF)**:
   - **Procedure**: Install the PSF Generator plugin from PSF Generator. Navigate to `Plugins -> PSF Generator -> Born & Wolf 3D Optical Model`. Set the acquisition parameters according to the imaging setup and run the generator. Save the PSF as a .tiff file.
   - **Rationale**: The PSF represents how a point source of light is perceived through the optical system. Generating a specific PSF for your setup allows for accurate

deconvolution, which enhances image resolution by reversing the blurring effects of the imaging system.

2. **Install the Deconvolution Plugin**:
   - **Procedure**: Download and install the DeconvolutionLab2 plugin from DeconvolutionLab2.
   - **Rationale**: DeconvolutionLab2 performs the deconvolution process, which is critical for improving image clarity and detail.
3. **Load the Fiji Macro for Deconvolution**:
   - **Procedure**: Load the Fiji macro `DeconLab.ijm` (different code file). Navigate to `Plugins -> Macros -> Startup Macros`. In the new window, go to `File -> Open` and select `DeconLab.ijm`.
   - **Rationale**: The macro automates the deconvolution process across all frames, ensuring consistency and efficiency.
4. **Edit Macro Paths**:
   - **Procedure**: Open the `DeconLab.ijm` macro file and edit the paths to point to your image frames, the generated PSF file, and the output directory.
   - **Rationale**: Correctly specifying the file paths ensures that the macro can access the necessary inputs and save the outputs appropriately.
5. **Adapt Loop Limits**:
   - **Procedure**: Modify the loop limits within the macro to match the number of frames in your time-lapse series.
   - **Rationale**: Ensuring the loop limits match the number of frames ensures that the deconvolution process is applied to all frames in the series.
6. **Run the Deconvolution Macro**:
   - **Procedure**: Click `Run` to execute the deconvolution process on the entire image series.
   - **Rationale**: Running the macro applies the deconvolution algorithm to all frames, enhancing image quality and preparing the data for further analysis.

#### Supplementary Note 2. Masking

The first step is masking the gel to avoid unwanted contribution from the outer background region.

**File name** `contractingActomyosinAutoMask`*.m*

##### Function Definition

```
[fileOut, sizeX, sizeY, percAreaMasked] =
contractingActomyosinAutoMask(frameIdx, sortedFrames, dirIn, dirOut)
```

- **Outputs (`fileOut`, `sizeX`, `sizeY`, `percAreaMasked`)**:

- `fileOut`: Output file name or path after processing.
- `sizeX`, `sizeY`: Dimensions of the processed image.
- `percAreaMasked`: Percentage of the image area that is masked or altered.

- **Inputs (`frameIdx`, `sortedFrames`, `dirIn`, `dirOut`)**:
  The function processes each frame (`frameIdx`) of images related to actomyosin contraction, analyzing and masking certain areas based on specified criteria.

  - `frameIdx`: Index of the frame to be processed.
  - `sortedFrames`: A collection or list of sorted frame names or identifiers.
  - `dirIn`: Directory path where input images are located.
  - `dirOut`: Directory path where output images will be saved.

**Constants (in the mask file)**

```
frameAccumulationCnt = 80; % Number of frames from which to accumulate
information.

percForAccumulation = 95; % Percentile value used for accumulation.

marginFreeBorder = 10; % Minimum distance from the border where no objects
are allowed.
```

- `frameAccumulationCnt`: Determines how many frames are considered for accumulation, for statistical operations.
- `percForAccumulation`: Sets the percentile value for accumulation, possibly used for thresholding or determining significant features.
- `marginFreeBorder`: Defines a buffer zone around the image border where processing might be restricted.

**Initialization**

```
file = sortedFrames(frameIdx);
pathToFile = [strcat(dirIn, file)];
image = imread(char(pathToFile));
[sizeX, sizeY, chanCnt] = size(image);
```

- `file`: Retrieves the filename from `sortedFrames` based on `frameIdx`.
- `pathToFile`: Constructs the full path to the input image file.
- `image`: Reads the image specified by `pathToFile` using `imread`.
- `[sizeX, sizeY, chanCnt]`: Retrieves the dimensions (`sizeX`, `sizeY`) and number of channels (`chanCnt`) of the image.

#### Supplementary Note 3. Main PIV file

**File name for continuous flow** `contractingActomyosinMain2.m`

Here is a detailed explanation of the `contractingActomyosinMain2.m` file, breaking down its structure and functionality. Below MATLAB script processes and analyses a set of `.tif` files, related to actomyosin contraction experiments. It performs various steps such as clearing previous data, setting directories, managing temporary files, copying batches of files for analysis, and running the PIV (Particle Image Velocimetry) analysis.

**Key Components**

1. **Clearing Previous Data:**

```
clc; clear all
```

This line clears the command window and all variables to ensure a fresh start for the script.

2. **Setting Directories and Parameters:**

```
dirIn          = 'D:/Work/BGU_Isreal/PIV/Gels_Analysis/...';
dirOut         = '/tmp/out/';
dirTmp         = '/tmp/tmp/';
framesDelta    =  300;
framesDeltaInnerStep = 1;
fileExtension = '.tif';
```

These lines define the input, output, and temporary directories along with other parameters like the frame interval (`framesDelta`) and file extension. `framesDelta` is the cross-correlation factor (most of the times 300 gives best result). `framesDeltaInnerStep(`$\Delta t$`)` can be changed as according to the dynamic flow.

3. **Checking Directory Conflicts:**

```
if (strcmp(dirTmp, dirIn) || strcmp(dirTmp, dirOut))
    disp("Warning: tmp folder needs to be separate from input and output
folders.");
end
```

This ensures the temporary directory is different from the input and output directories to avoid conflicts.

4. **Creating and Cleaning Directories:**

```
if exist(dirTmp, 'dir')
    rmdir(dirTmp, 's');
end
if exist(dirOut, 'dir')
    rmdir(dirOut, 's');
end
mkdir(dirTmp);
```

This part checks if the temporary and output directories exist, deletes them if they do, and then creates fresh directories.

5. **Reading and Sorting Files:**

```
directoryContents = dir([dirIn, filesep, ['*' fileExtension]]);
filenames = {directoryContents.name};
filenames = sort_nat(filenames);
```

This reads all files with the specified extension from the input directory, stores their names, and sorts them naturally using the `sort_nat` function (which must be installed separately).

6. **Processing Files in Batches:**

```
amount = length(filenames);
for startIdx = 1:amount - 1
    clearFolder(dirTmp, fileExtension);
    approxVelocity = getApproxRadiusVelocity(startIdx, framesDelta,
filenames, dirIn, dirOut);
    maskReady = false;
    for i = 1:framesDeltaInnerStep:framesDelta
        idx = startIdx + i;
        if (idx <= amount)
            file = filenames(idx);
            pathToFile = strcat(dirIn, file);
            copyfile(char(pathToFile), dirTmp);
            if (maskReady == false)
                [maskFile, sizeX, sizeY, percAreaMasked] =
contractingActomyosinAutoMask(idx, filenames, dirIn, dirOut);
                maskReady = true;
            end
        end
    end
    [maskiererx , maskierery] = loadMask(maskFile, framesDelta);
    PIVanalysis(dirTmp, dirOut, framesDelta, approxVelocity, sizeX,
sizeY, percAreaMasked, maskiererx, maskierery);
```

```
end
```

This loop processes files in batches. It starts by clearing the temporary directory, calculates an approximate velocity, copies files to the temporary directory, and checks if a mask is ready. Then, it runs the PIV analysis on the batch of files.

7. **Helper Functions:**
   - **`getApproxRadiusVelocity`**: Computes an approximate velocity based on the radius of the masked area.
   - **`contractingActomyosinAutoMask`**: Generates a mask for the image files to identify areas of interest.
   - **`loadMask`**: Loads and processes the mask for PIV analysis.
   - **`PIVanalysis`**: Performs the PIV analysis on the batch of files.

**Main Functions Explanation**

- **`getApproxRadiusVelocity`**: This function calculates the velocity by examining two points in time, separated by `framesDelta`. It uses the `contractingActomyosinAutoMask` function to get the mask and compute the area, then determines the radius and velocity.
- **`contractingActomyosinAutoMask`**: Generates a mask for the image, which is used to isolate the region of interest for analysis.
- **`loadMask`**: Loads the mask file and prepares it for PIV analysis.
- **`PIVanalysis`**: Performs Particle Image Velocimetry analysis on the batch of files in the temporary directory, using the prepared mask and other parameters.

**Standard PIV Settings.**

The section of code provided sets up various parameters for Particle Image Velocimetry (PIV) analysis. These parameters configure different preprocessing steps, PIV settings, and options that influence the quality and accuracy of the analysis.

#### Detailed Explanation

1. **Initialization of Parameters Cell Array:**

```
s = cell(25,2);
```

Creates a cell array `s` with 25 rows and 2 columns. Each row corresponds to a different parameter, where the first column (`s{:,1}`) holds the parameter name and the second column (`s{:,2}`) holds the parameter value.

2. **Setting Parameters:**

```
% Parameter                          % Setting
s{1,1}  = 'Auto contrast stretch';  s{1,2}  = 1;
s{2,1}  = 'Video frame selection';  s{2,2}  = [];
s{3,1}  = 'Background image A';     s{3,2}  = [];
s{4,1}  = 'Background image B';     s{4,2}  = [];
s{5,1}  = 'CLAHE';                  s{5,2}  = 1;
s{6,1}  = 'Highpass';               s{6,2}  = 0;
s{7,1}  = 'Clipping';               s{7,2}  = 0;
s{8,1}  = 'CLAHE size';             s{8,2}  = 64;
s{9,1}  = 'Highpass size';          s{9,2}  = 15;
s{10,1} = 'Wiener';                 s{10,2} = 0;
s{11,1} = 'Wiener size';            s{11,2} = 11;
s{12,1} = 'ROI';                    s{12,2} = '';
s{13,1} = 'maskiererx';             s{13,2} = maskListA;
s{14,1} = 'maskierery';             s{14,2} = maskListB;
s{15,1} = 'Int. area 1';            s{15,2} = 128;
s{16,1} = 'Step size 1';            s{16,2} = 64;
s{17,1} = 'Subpix. finder';         s{17,2} = 1;
s{18,1} = 'Nr. of passes';          s{18,2} = 4;
s{19,1} = 'Int. area 2';            s{19,2} = 64;
s{20,1} = 'Int. area 3';            s{20,2} = 32;
s{21,1} = 'Int. area 4';            s{21,2} = 16;
s{22,1} = 'mask_auto';              s{22,2} = 0;
s{23,1} = 'Window deformation';     s{23,2} = "linear";
s{24,1} = 'Repeated Correlation';   s{24,2} = 0;
s{25,1} = 'Use padding';            s{25,2} = 1;
```

Each `s{i,1}` is a parameter name, and `s{i,2}` is its corresponding setting.

- **Parameter Explanations:**
  - **Auto contrast stretch**: Enhances contrast, important for 16-bit images.
  - **Video frame selection**: Used if input is a video.
  - **Background image A/B**: Paths to background images for subtraction.
  - **CLAHE**: Adaptive histogram equalization for contrast enhancement.
  - **Highpass**: Highpass filter to remove low-frequency noise.
  - **Clipping**: Removes extreme values to reduce noise.
  - **CLAHE size**: Size of the CLAHE window.
  - **Highpass size**: Size of the highpass filter.
  - **Wiener**: Adaptive denoise filter.
  - **Wiener size**: Size of the Wiener filter.
  - **ROI**: Region of Interest for analysis.
  - **maskiererx/maskierery**: Mask coordinates.
  - **Int. area 1/2/3/4**: Sizes of interrogation windows for multiple passes.

- **Step size 1**: Step size for the first pass.
- **Subpix. finder**: Subpixel displacement finder method.
- **Nr. of passes**: Number of passes for PIV analysis.
- **mask_auto**: Automatic masking.
- **Window deformation**: Method for window deformation ('linear' or 'spline').
- **Repeated Correlation**: Whether to repeat correlation and average results.
- **Use padding**: Padding to improve PIV accuracy.

#### Supplementary Note 4. Interrogation Window adaptation and step size

```
% Adapt the interrogation areas to the size of the mask.
s{15,2} = radius / 1;           % Int. Area 1.
```

- This line sets the first interrogation area size (`s{15,2}`) to be equal to the `radius` of the mask.
- The division by `1` means no scaling is applied here, so the interrogation area is directly set to the size of the mask's radius.

```
s{16,2} = floor(s{15,2} / 2);  % Step size.
```

- This line sets the step size (`s{16,2}`) to half the size of the first interrogation area.
- The `floor` function ensures that the result is an integer by rounding down to the nearest whole number.

```
s{19,2} = s{16,2};              % Int. Area 2.
```

- This line sets the second interrogation area (`s{19,2}`) to be the same as the step size determined in the previous line.
- This means the second interrogation area is half the size of the first interrogation area.

```
s{20,2} = floor(s{19,2} / 2);  % Int. Area 3.
```

- This line sets the third interrogation area (`s{20,2}`) to half the size of the second interrogation area.
- Again, the `floor` function is used to ensure the value is an integer.

```
s{21,2} = floor(s{20,2} / 1);  % Int. Area 4.
```

- This line sets the fourth interrogation area (`s{21,2}`) to be the same size as the third interrogation area.

- The division by `1` means no further scaling is applied here, so the size remains the same as the third interrogation area.

#### Supplementary Note 5. Strain analysis

1. Strain calculation: The part of MATLAB code to calculate the strain is shown below.

```
dx = x(2) - x(1);
dy = y(2) - y(1);
for i = 2:height(u_x)-1
    for j = 2:width(u_y)-1
        u_xx(i,j) = (u_x(i,j+1)-u_x(i,j-1))/(2*dx);
        u_yy(i,j) = (u_y(i+1,j)-u_y(i-1,j))/(2*dy);
        u_xy(i,j) = 1/2*((u_x(i+1,j)-u_x(i-1,j))/(2*dy)
        +(u_y(i,j+1)-u_y(i,j-1))/(2*dx));
    end
end
```

Here `u_xx`, `u_yy` and `u_xy` are the components of strain $\varepsilon_{xx}$, $\varepsilon_{yy}$ and $\varepsilon_{xy}$ respectively.

2. Cartesian to polar transformation

Location of the geometric center (centroid) of gel:

```
XCM_new = mean(x,'all','omitnan');
YCM_new = mean(y,'all','omitnan');
```

Calculation of polar angle:

```
x = x - XCM_new;
y = y - YCM_new;
r = sqrt(x.^2+y.^2);
phi = atan(y,x);
```

Displacement from cartesian to polar components:

```
u_r = u_x.*cos(phi) + u_y.*sin(phi);
u_phi = -u_x.*sin(phi) + u_y.*cos(phi);
```

Strain from cartesian to polar components:

```
u_rr = u_xx.*cos(phi).^2 + u_yy.*sin(phi).^2 +
2*u_xy.*sin(phi).*cos(phi);
u_phiphi = u_xx.*sin(phi).^2 + u_yy.*cos(phi).^2 -
2*u_xy.*sin(phi).*cos(phi);
u_rphi = -(u_xx - u_yy).*sin(phi).*cos(phi) + u_xy.*(cos(phi).^2 +
sin(phi).^2);
```

Here `u_rr`, `u_phiphi` and `u_rphi` are the components of strain $\varepsilon_{rr}$, $\varepsilon_{\phi\phi}$ and $\varepsilon_{r\phi}$ respectively.

3. The following lines of MATLAB code is used to generate a 2D heatmap of bulk area strain:

```
U = u_xx+u_yy;
Umin = min(U,[],'all');
Umax = max(U,[],'all');
U(U>0)=U(U>0)/Umax;
U(U<0)=-U(U<0)/Umin;
contourf(x,-y, U, 50, 'EdgeColor','none')
hold on
quiver(x,-y,u_x,-u_y,'r', 'ShowArrowHead','on')
plot(XCM_new,-YCM_new,'o','MarkerSize', 7, 'MarkerEdgeColor', 'r',
'LineWidth', 2)
hold off
set(gca, 'Color', 'k')
colorbar
clim([-1 1])
axis square
```

4. Calculation of eigenvalues and eigenvectors of the strain

```
EigenvectorxT = NaN(height(u_xx), width(u_xx));
      EigenvectoryT = NaN(height(u_xx), width(u_xx));
      Eigenvectorxt = NaN(height(u_xx), width(u_xx));
      Eigenvectoryt = NaN(height(u_xx), width(u_xx));
      EigenvalueT = NaN(height(u_xx), width(u_xx));
      Eigenvaluet = NaN(height(u_xx), width(u_xx));
      for i = 1:size(u_xx,1)
            for j = 1:size(u_xx,2)
            A = [u_xx(i,j) u_xy(i,j); u_xy(i,j) u_yy(i,j)];
            [V,D] = eig(A);
            Eigvec = reshape(V, [4,1]);
            Eigval = reshape(D, [4,1]);
            [T,I] = max(abs(Eigval), [], "all");
            Tmax= Eigval(I);
            dd = Eigval;
            dd = abs(dd);
            dd(dd==0) = inf;
            [t,J] = min(dd, [], "all");
            Tmin = Eigval(J);
            if I==1
                  EigenvectorxT(i, j) = Eigvec(I);
              EigenvectoryT(i, j) = Eigvec(I+1);
              Eigenvectorxt(i, j) = Eigvec(I+2);
```

```
            Eigenvectoryt(i, j) = Eigvec(I+3);
        elseif I==4
            EigenvectorxT(i, j) = Eigvec(I-1);
            EigenvectoryT(i, j) = Eigvec(I);
            Eigenvectorxt(i, j) = Eigvec(I-3);
            Eigenvectoryt(i, j) = Eigvec(I-2);
        end
        EigenvalueT(i,j) = Tmax;
        Eigenvaluet(i,j) = Tmin;
        end
end
```

The 2D heatmaps of eigenvalues (Figure. 8) can be generated in the same way as 2D area strain plots, by substituting `EigenvalueT` in the definition of `U` in the code and substituting the displacement vectors with eigenvectors, i.e. '`u_x, u_y`' with '`EigenvectorxT, EigenvectoryT`'

### Supplementary Figures

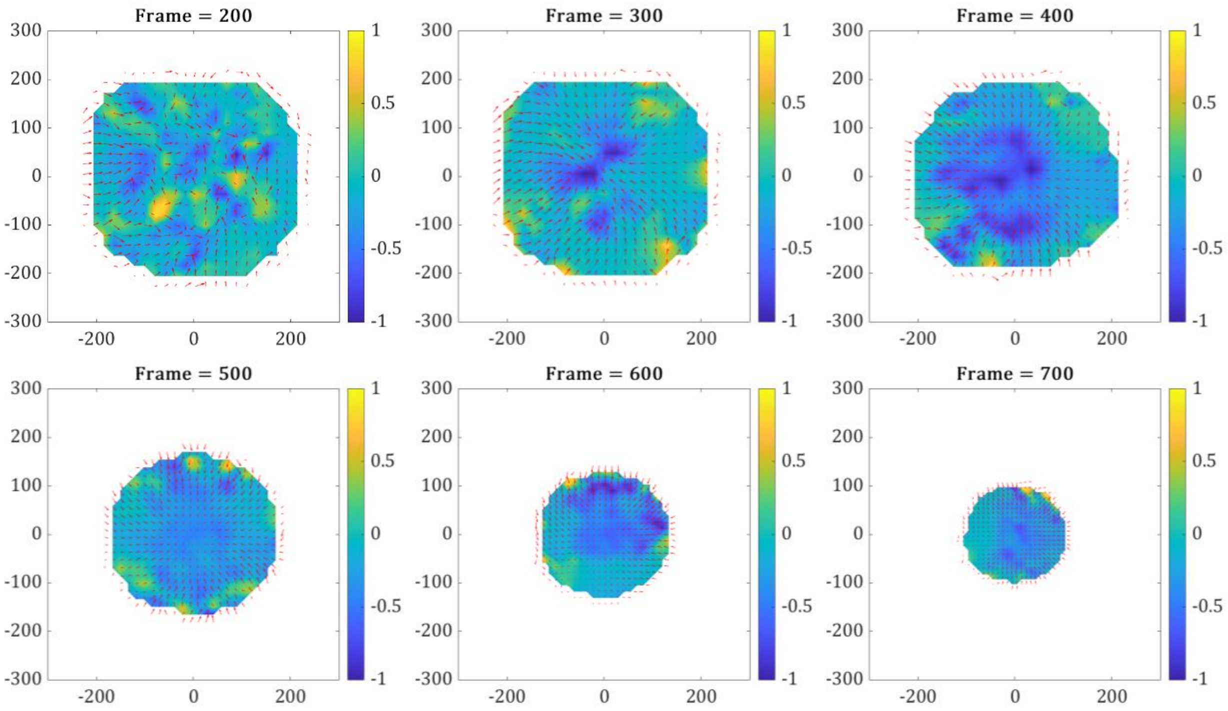


**Figure S1. Time evolution of area strain for the gel disc shown in Fig. 1.** Different snapshots of the trace of the strain (color) and displacement vectors (red arrows) obtained from PIV analysis of different frames of the contracting gel. These frames correspond to the single representative frame of the gel shown in Fig. 7. After an initial period when contraction is building up, the contraction proceeds primarily isotopically in the inner region (blue), with some stretched regions near the boundary (yellow).

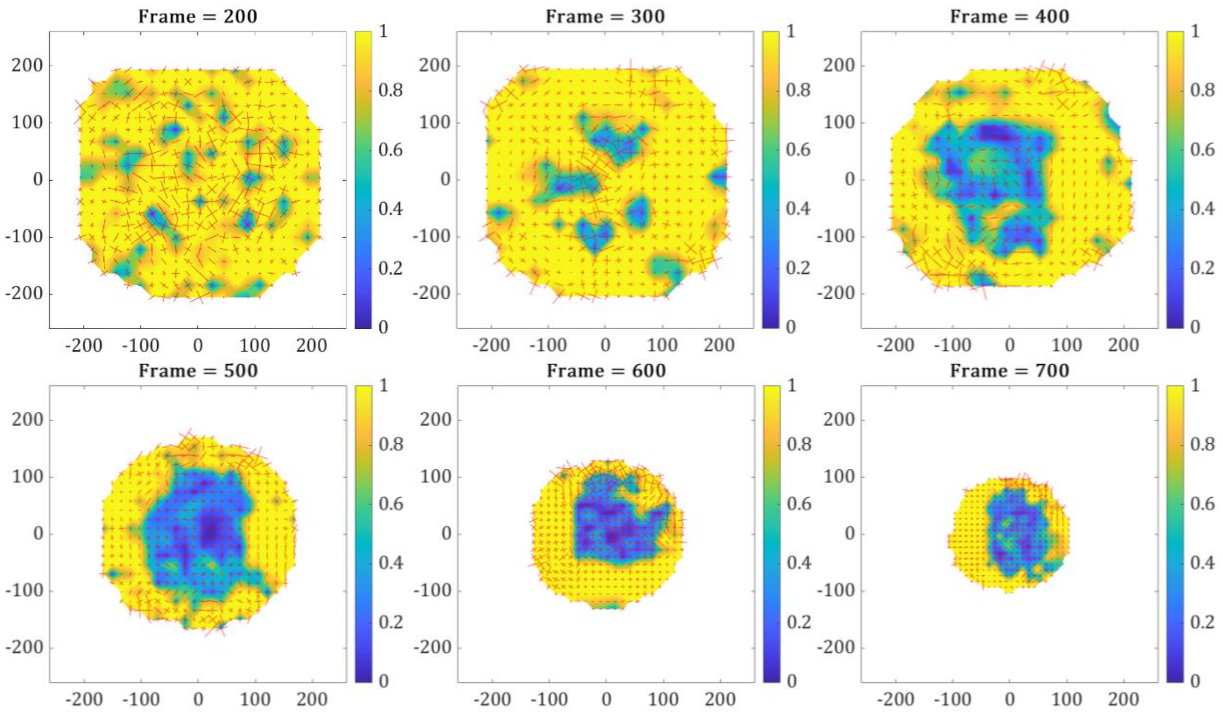


**Figure S2. Time evolution of 2d maps of normalized difference of eigenvalues for the gel depicted in Fig. 1.** The color map shows the normalized difference of the two strain eigenvalues for several representative frames from PIV of the circular gel, corresponding to the representation shown in Fig. 8 in the main text. Blue regions correspond to isotropic contraction, whereas yellow indicates unequal contraction along radial and azimuthal directions The vector map shows the two orthogonal eigenvectors scaled by the magnitude of its corresponding eigenvalue.

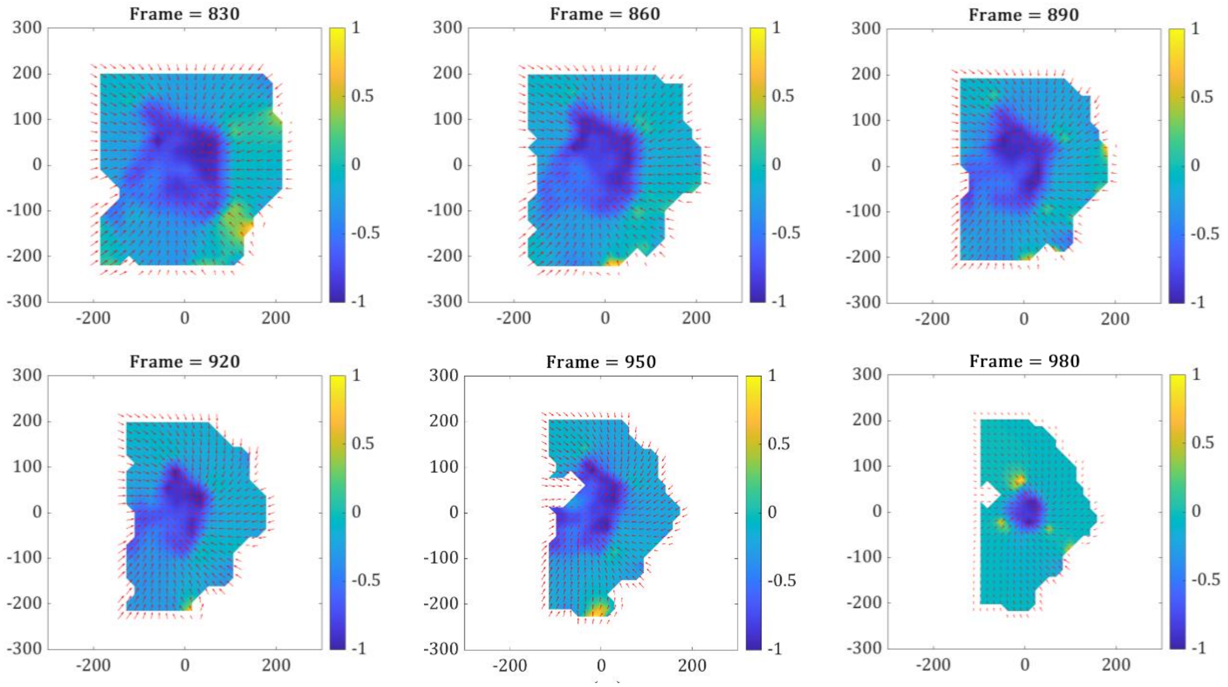


**Figure S3. Time evolution of area strain for irregular gel depicted in Fig. 11.** Different snapshots of the trace of the strain (color) and displacement vectors (red arrows) obtained from PIV analysis of different frames of a contracting non-circular gel shown in Fig. 11 in the main text. Strongly blue regions represent an isotropically contracting inner core region surrounded by patches of stretched regions (yellow).

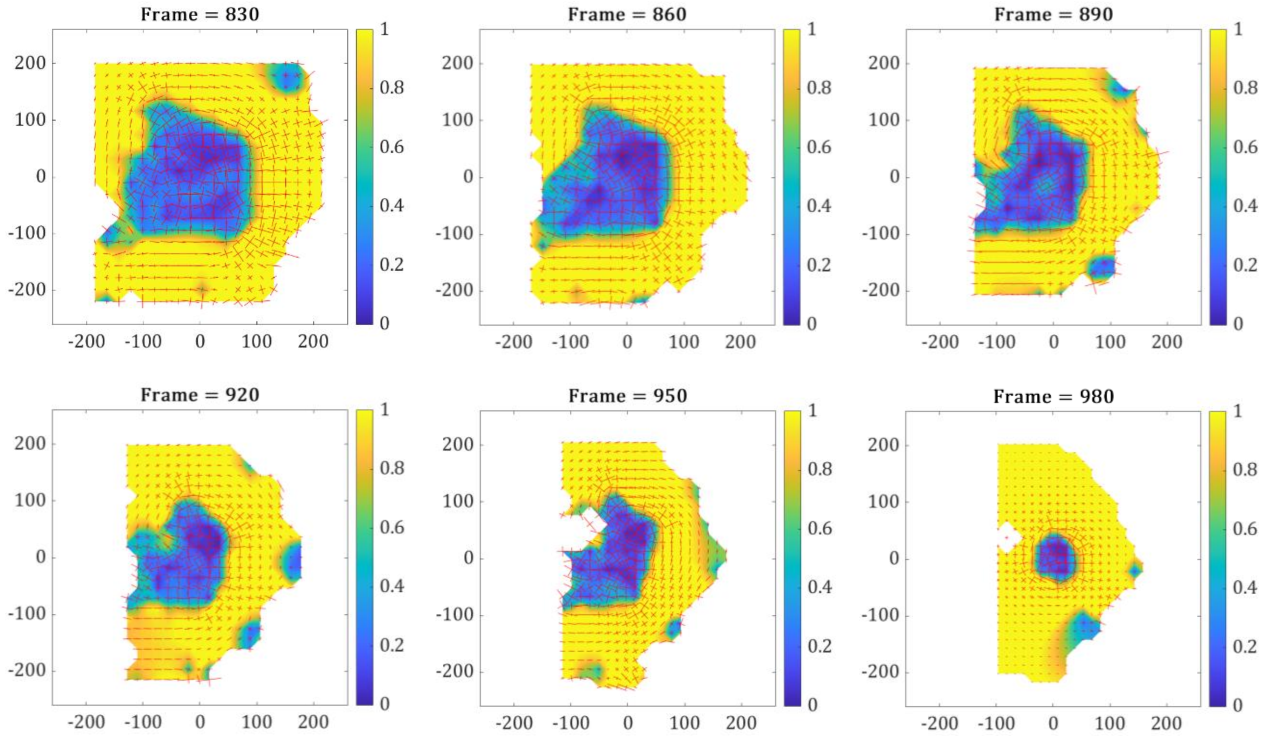


**Figure S4. Time evolution of 2d maps of normalized difference of eigenvalues for the gel shown in Fig. 11.** The color map shows the normalized difference of the two strain eigenvalues for several representative frames from PIV of the non-circular gel shown in Fig. 11 of the main text. In all frames, there is an isotropic inner region (blue) where the two eigenvalues are equal, whereas the surrounding region shows strong strain anisotropy (yellow). This anisotropy is reflected in the vector map, which shows orthogonal eigenvectors, scaled by the corresponding eigenvalue. In almost all cases seen, the dominant eigenvector is azimuthal.
